# Development of 50 Nanometer Diameter Genetically Encoded Multimeric Nanoparticles as Tools to Probe the Physical Properties of the Cytoplasm

**DOI:** 10.1101/2024.07.05.602291

**Authors:** Cindy M. Hernandez, David C. Duran-Chaparro, Trevor van Eeuwen, Michael P. Rout, Liam J. Holt

## Abstract

The mechanisms that regulate the physical properties of the cell interior remain poorly understood, especially at the mesoscale. Many crucial macromolecules and molecular assemblies such as ribosomes, RNA polymerase, and biomolecular condensates span the mesoscale size range, and changes in mesoscale physical properties have been suggested to be crucial for both normal physiology and disease. Therefore, we need better tools to study the cellular environment at this scale. Physical properties of the cell can be inferred through analysis of the motion of tracer nanoparticles, an approach called nanorheology. This requires the introduction of nanoparticles to cells, which can be labor intensive and slow. *Genetically Encoded Multimeric nanoparticles* (GEMs) were recently developed to address this limitation. GEMs consist of scaffold proteins fused to fluorescent tags that self-assemble into bright and stable nanoparticles of defined geometry. However, extremely sensitive microscopes were required to track previous (40nm-GEMs). Here, we describe the development and characterization of 50 nm diameter GEMs (50nm-GEMs) that are brighter and probe a slightly longer length scale. 50nm-GEMs will make high-throughput nanorheology accessible to a broader range of researchers and reveal new insights into the biophysical properties of cells.

## Introduction

Eukaryotic cells are crowded with macromolecules, membrane-bound organelles, and a dynamic cytoskeleton^1–5–6^. The diffusion of particles within this complex environment can be decreased by higher viscosity, macromolecular crowding and elastic confinement, but increased by metabolic activity, polymerization and depolymerization (especially of cytoskeletal networks), and the activities of molecular motors, such as myosins, kinesins, and helicases.

It is of particular importance to understand the physical properties of the intracellular environment at the mesoscale (~10 to 1000 nanometers diameter) because a large fraction of the space in the cell is taken up by particles of this size (e.g. nucleosomes are ~10 nm, ribosomes are ~30 nm, mRNA ribonucleoproteins complexes are ~100 nm). Most monomeric proteins are in the nanometer (1-5 nm) size range^3,4,5^. However, a large proportion of a cell’s proteins assemble into mesoscale complexes, including macromolecular complexes that drive key processes such as transcription, translation and DNA replication^6,7^. Furthermore, recent experiments suggest that at least 18% of proteins in the *Xenopus laevis* cytoplasm are organized into mesoscale biomolecular condensates of predominantly 100 nanometer diameter^8^. Therefore, we need better technologies and approaches to understand how the intracellular environment influences mesoscale particles.

Some of the most successful characterizations of the cell interior to date have employed **nanorheology**^7^, which is the inference of biophysical properties from observations of the motion of tracer particles^8^. A major challenge is the shortage of methods to introduce probes for nanorheology into cells. Approaches such as microinjection of tracer particles into cells can damage cells and dilute the cytoplasm; furthermore, these approaches are labor intensive and low-throughput.

To address these challenges, our lab has previously reported the development of Genetically Encoded Multimeric (GEMs) nanoparticles of 20 and 40 nanometer diameter (20- and 40nm-GEMs respectively)^9,10^. We based these probes on scaffold proteins that self-assemble into icosahedral structures of precise geometry. We fused these scaffold proteins to fluorescent proteins yielding bright, stable nanoparticles. We can use different scaffolds to make GEMs of different sizes, allowing us to probe the physical properties of the cell at different length-scales. GEMs are constantly expressed, solving all of the problems mentioned above: no dilution occurs; the cortex and membrane are not disrupted; and every cell contains nanoparticles, enabling acquisition of data from thousands of cells. Moreover, GEMs are well tolerated ^11,12^, allowing long-term studies and high content genetic screens. GEMs have a well-defined geometry and size allowing precise physical interpretation of their motion. They are bioorthogonal, and therefore unlikely to be subject to regulated interactions with the cell. GEMs are easy to use: no microinjection or laborious sample preparation is required, enabling the nanorheological characterization of many species for the first time. For example, it is impractical to micro-inject microorganisms that have cell walls, including plants or yeasts, but GEMs can be easily introduced into these organisms ^12–15^.

Genetically encoded multimeric nanoparticles (GEMs) have been successfully applied across a wide range of biological systems, from prokaryotes^10^ to multicellular eukaryotes^11^, to probe the physical properties of the intracellular environment. In *Escherichia coli*, GEMs have been used to reveal cytoplasmic organization and exclusion of ribosome-sized particles from the nucleoid^10^. In *Saccharomyces cerevisiae*, GEMs have provided insights into physical alterations associated with cellular aging^12^, the physical impacts of compression^13^, the link between crowding, activity and condensate function^14,15^, and the effects of nutrient deprivation on cytoplasmic organization^16^. In *Schizosaccharomyces pombe*, GEMs have provided key insights into nuclear size scaling^17^. Furthermore, GEM-based approaches have contributed to understanding polysome collapse and RNA-associated phase behavior during cellular stress^18^. GEMs have also been employed in *Caenorhabditis elegans* to study the biophysical properties of cells in animal tissues^11^. In mammalian cells they have been used to study the deformability of the nucleus^19^ and physical properties of the cytoplasm^20,21^ and to examine the manipulation of nuclear physical properties by viruses^22^ Together, these studies demonstrate the versatility of GEMs as a tool to investigate intracellular physical states across diverse organisms and biological contexts.

The development of 40nm-GEMs has accelerated discovery. However, expansion of the available probes within the mesoscale range would provide deeper insights into the biophysical properties of the cell interior. Furthermore, previous GEMs are challenging to use at sufficient temporal resolution (30-100 Hz) because highly sensitive microscopes are required to achieve sufficient signal at these rapid frame-rates. Thus, in this study, we expand our nanorheology probe toolkit by developing a 50 nm diameter GEM using the *Quasibacillus thermotolerans* encapsulin^16^ as a scaffold. These “50nm-GEMs” are brighter, easier to use, provide longer tracks and probe the biophysical properties of the cell at the 50 nm length-scale.

## Results

### 50 nm-GEMs are robustly expressed and assembled and exhibit greater size and brightness than 40nm-GEMs in yeast *S. cerevisiae*

We engineered a 50 nm diameter GEM (50nm-GEM) by fusing the Green Fluorescent Protein (GFP^23^) specifically Super Folder GFP (sfGFP^24^), or a Sapphire^25^ variant differing from the canonical sequence by two conservative substitutions (H231T and M233I) to the C-terminus of an encapsulin cage protein from *Quasibacillus thermotolerans*. Prior structural work^26^ found that this encapsulin self-assembles into a 240-subunit icosahedral compartment of ~42 nm in diameter. Because each subunit is fused to a fluorescent protein, we predicted that the assembled GEM would reach an effective diameter of ~50 nm due to the surface “cloud” of fluorophore molecules (**Figure 1A**). This length-scale is at a critical threshold at which the mesoscale environment is thought to become glassy and depend upon metabolic activity for particle motion^27^.

**Figure 1.**
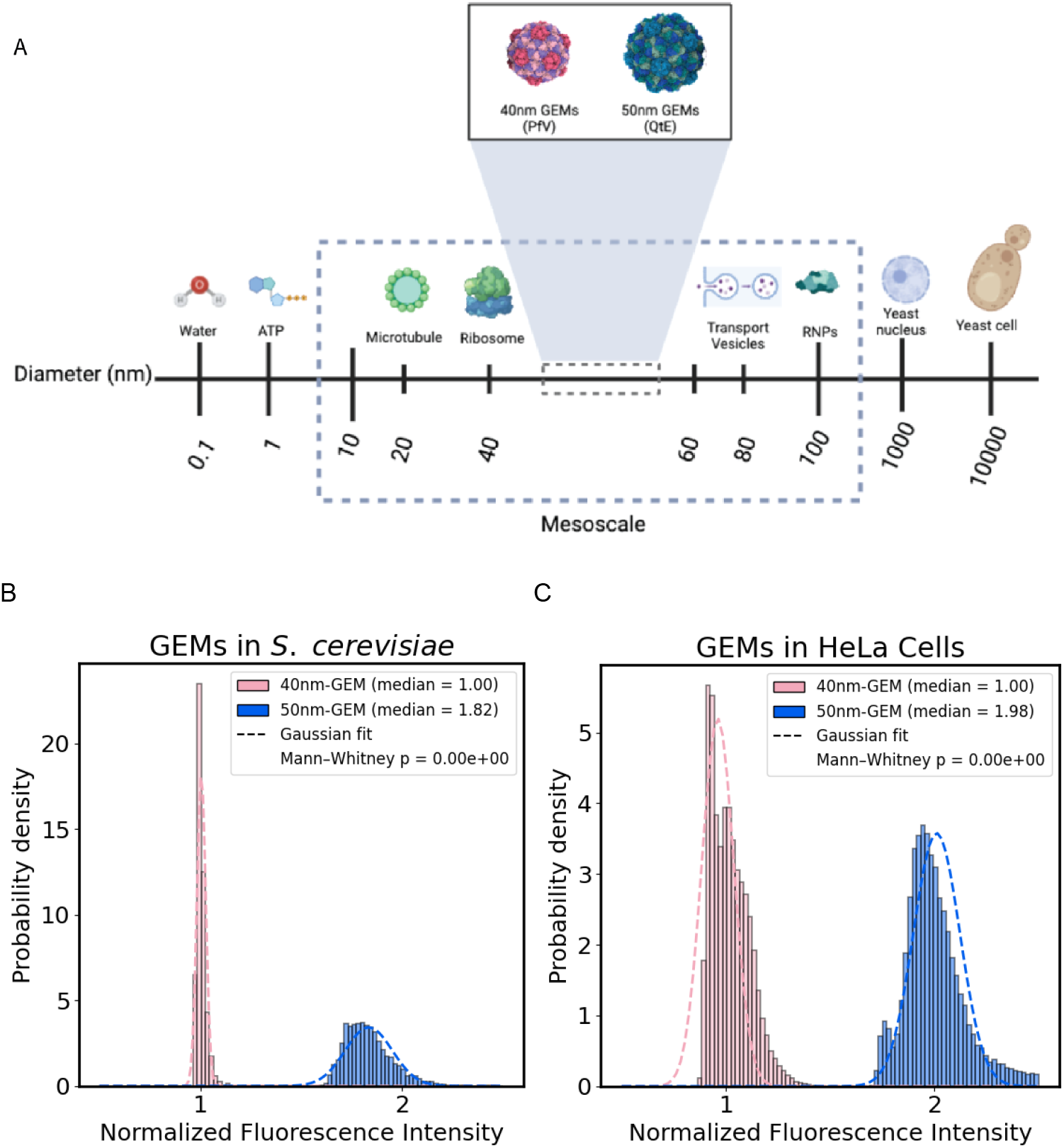
50nm-GEMs are brighter than 40nm-GEMs. **(A)** Schematic showing relative sizes of key biomolecules, organelles and organisms compared to GEM nanoparticles. **(B)** Fluorescence intensity distributions of GEMs in yeast (*S. cerevisiae)* and mammalian (HeLa) cells. Histograms show the normalized fluorescence intensity of 40nm-GEMs (pink) and 50nm-GEMs (blue) in yeast (left) and mammalian (right) cells. Dashed lines are Gaussian fits. Mann–Whitney tests confirmed that 50nm-GEMs are significantly brighter than 40nm-GEMs ((*p* < 0.0001)). **Yeast data** include 24,594 tracked particles, with ≥300 cells per replicate analyzed across *N* = 3 independent biological replicates. Differences in median normalized fluorescence intensity were evaluated using a Mann–Whitney test (p < 0.0001). **Mammalian data** comprise 91,333 tracked particles, quantified from ≥50 cells per replicate across *N* = 3 biological replicates. Differences in median normalized fluorescence intensity were assessed using a Mann–Whitney test (p < 0.0001).

The *Q. thermotolerans* encapsulin scaffold of 50nm-GEMs self-assembles from 240-monomers, as opposed to the *Pyrococcus furiosus* scaffold of the 40nm-GEMs, which assembles from 180 monomers. Therefore, 50nm-GEMs should be substantially brighter because fluorophore number, and therefore brightness, scales with monomer count. To test this, we plotted the brightness of detected particles normalized to the median brightness of 40nm-GEMs. These distributions have narrow, well-defined peaks, suggesting that particle assembly is relatively uniform. Mammalian GEMs have slightly broader distributions of brightness. Overall, we found that 50nm-GEMs displayed approximately double the fluorescence intensity of 40nm-GEMs in both *S. cerevisiae* and HeLa cells (**Figure 1B,C**).

The necessity to image GEMs rapidly (30-100 Hz) limits us to two-dimensional (2D) imaging and also limits the signal to noise of our data. It can also be challenging to obtain cells with low numbers of GEM nanoparticles, leading to possible artifacts from simultaneous assignment of multiple GEMs as a single particle, or incorrect tracking where multiple GEMs are incorrectly assigned to a single track. To test the impact of these possible artifacts and optimize our analysis pipeline, we needed a dataset with a known ground-truth. Therefore, we generated synthetic movies of fluorescent particles **(Supplementary Figure 1)**. We simulated trajectories of particles undergoing three-dimensional Brownian diffusion with a uniform diffusion coefficient matched to the median diffusivities of GEMs **(Supplementary Figure 1A, left)**. To attempt to mimic our real data, images were then rendered by projecting a diffraction-limited Airy disk point spread function, with axial position on a pixel grid using and modulating emitter brightness and apparent size to mimic defocus **(Supplementary Figure 1A, right)**.. Noise levels were tuned to match the signal-to-noise ratio of the original movies **(Supplementary Figure 1A, center right)**. To capture the effects of limited focal volume in our 2D experimental imaging, particles entering a focal-plane slab were recorded and projected onto the imaging plane to reproduce experimental acquisition **(Supplementary Figure 1A, center left)**. The resulting stacks were saved as multi-frame TIFF files and processed through the same detection, tracking, and analysis pipeline used for experimental datasets **(Supplementary Figure 1A, right)**.

**Supplementary Figure 1.**
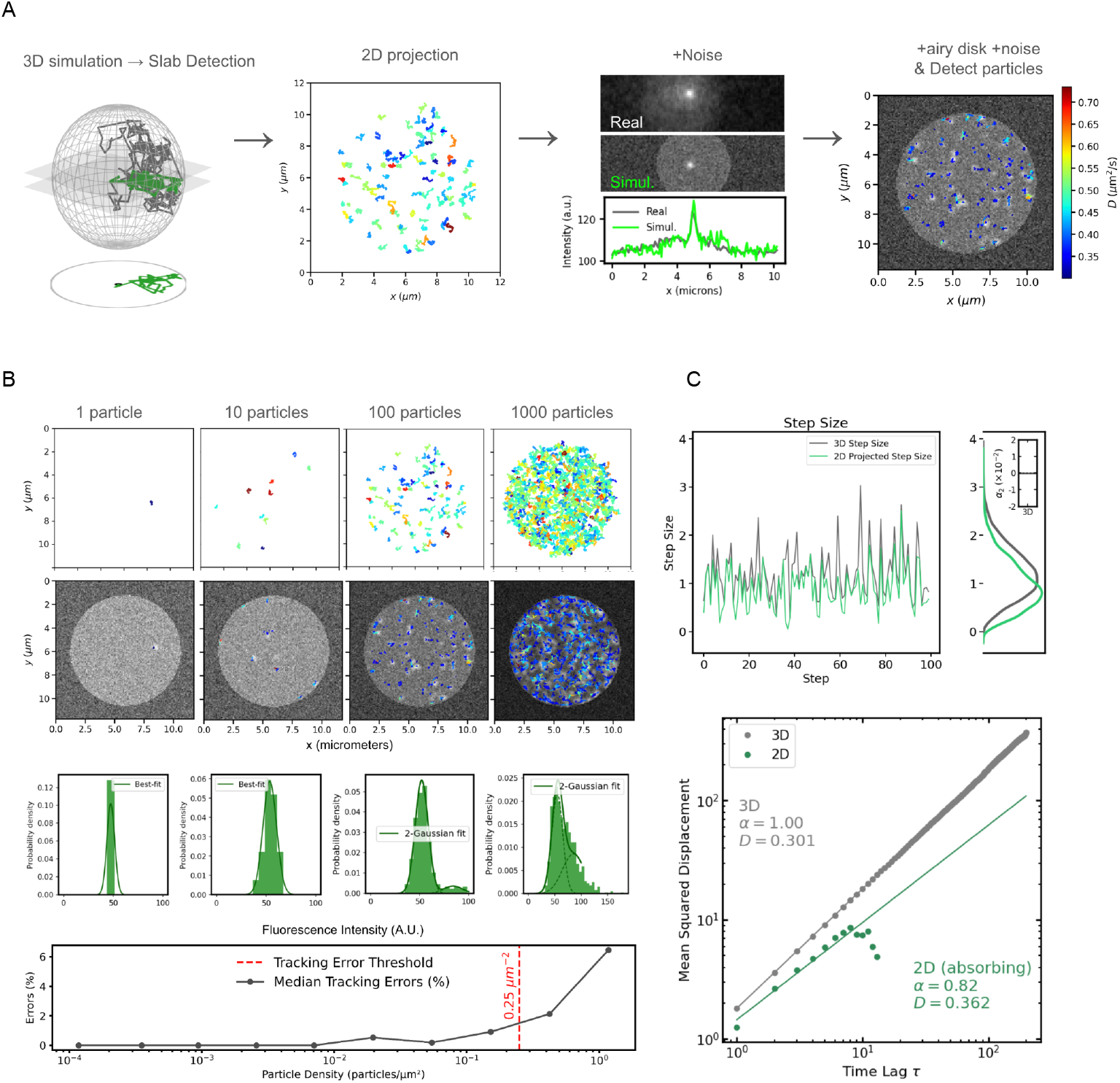
Particle density modulates tracking accuracy in simulated datasets. **(A) Left**. Schematic of particle motion within a bounded three-dimensional volume. Purple trajectories represent particles tracked throughout the full volume; green trajectories are sub-sections of tracks that can be detected within a thin optical section (shown below as a two-dimensional projection), mimicking fluorescence imaging in a single focal plane. **Center** Ground-truth two-dimensional trajectories illustrating diffusive motion across the field of view. Tracks are colored to distinguish different trajectories. **Right**. Simulated fluorescence microscopy image with detected particle tracks overlaid and colored by track. Movies were generated by convolving simulated trajectories with an Airy disk and incorporating defocus and imaging noise. Noise was simulated as described in the Methods section. The simulated movies were then analyzed using the same tracking pipeline that we applied to our experimental data. Each track has a different color. **(B) Top** Synthetic fluorescence images containing increasing particle densities (1, 10, 100, and 1000 particles). All particles undergo ideal Brownian motion with identical ground-truth diffusion properties. As density increases, spatial overlap and trajectory ambiguity progressively rise. Tracks are each assigned different colors. **Center**. Fluorescence intensity distributions corresponding to each density condition on top. Histograms were fit with a two-Gaussian mixture model representing detection of a single particle or several particles on top of each other at low particle densities, intensity distributions approximate Gaussian statistics, whereas higher densities produce broader, non-Gaussian distributions due to particle overlap fluorescence. **Bottom**. Tracking errors (particle identity swaps; see Methods) quantified as a function of particle density. The median tracking error rate (blue circles) remains minimal at low densities but increases sharply beyond a critical density. Shaded regions indicate the 95% confidence interval. The dashed red vertical line marks the tracking error threshold used throughout the study, defined as the density at which the slope of the error curve becomes significantly greater than zero. Experimental datasets were restricted to densities below this threshold to minimize identity confusion. **(C) Top Left**. Comparison of single-step displacements measured in three dimensions (purple) and after two-dimensional projection (green) for the same trajectory. Projection underestimates displacement magnitude by eliminating the axial component of motion. **(Top Right)** Probability density distributions of step sizes for full 3D trajectories (purple) and their 2D projections (green). Projection shifts the distribution toward smaller apparent displacements. *Inset:* Non-Gaussian parameter increases for projected trajectories, indicating stronger deviations from Gaussian statistics introduced by dimensional reduction and finite observation depth. **(Bottom)** Mean squared displacement (MSD) versus time lag (τ) for true 3D trajectories and corresponding 2D absorbing projections, in which trajectories terminate once particles exit the focal slab. Even though simulated particles move with strictly Brownian motion (α = 1), the projected trajectories exhibit apparent subdiffusive scaling (α < 1) arising solely from observation geometry. Only the first ten time lags are shown to minimize contributions from long-lag statistical noise.

### Isolation and Cryo-EM Single Particle Analysis of In vivo–assembled GEMs in *S. cerevisiae*

To understand the size and nature of 40nm-GEMs and 50nm-GEMs, we sought to leverage our previously developed methods for single step native affinity purification and structural characterization by cryo-electron microscopy single particle analysis (cryo-EM SPA) of large macromolecular complexes^28^. We first set out to determine whether we could isolate *in vivo* assembled GEMs expressed in *S. cerevisiae* by adapting our cryogenic milling and GFP affinity isolation protocol through α-GFP affinity isolation^29,30^ (**Figure 2A**). Both 40nm-GEMs and 50nm-GEMs were successfully isolated with high purity using polyclonal GFP antibodies (**Supplementary Figure 2a)**, but this strategy is not suitable for native elution and structural analysis. Since our goal was to analyze the *in vivo* assembled GEMs with the surface cloud of GFP molecules, we fused a FLAG-tag sequence to the C-terminus of the fluorescent tag^31^ to enable native elution. 40nm-GEMs and 50nm-GEMs purified by affinity capture and native elution by FLAG-peptide exhibited the same molecular weight and higher purity than those isolated by α-GFP affinity isolation (**Supplementary Figure 2a**) and were suitable for structural characterization. Negative stain transmission electron microscopy (TEM) of purified 50nm-GEMs revealed whole GEM particles consistent in size and shape with previous structural studies of the *Quasibacillus thermotolerans* encapsulin-based iron storage compartment^31^. Cryo-TEM imaging of both 50nm-GEMs and 40nm-GEMs showed large particles (**Supplementary Figure 2c)** and 2D class averages showed intact GEMs with clear features (**Supplementary Figure 2d**). After filtering particles in good 2D class averages, we obtained 3D reconstructions of 2.85 and 4.13 angstroms for the 50nm-GEMs and 40nm-GEMs, respectively (**Figure 2B,C left, Supplementary Figure 3**). These reconstructions were generally similar to previously characterized recombinant assemblies of the *Quasibacillus thermotolerans*^*31*^ *and Pyrococcus furiosus* encapsulins. The high-resolution GEM reconstructions, while providing a detailed structure of the scaffold, lacked significant density for the attached GFP proteins, likely due to their flexible attachment to the nanoparticle scaffold. To obtain information about the full hydrodynamic radius of the GEMs, we therefore repeated the refinement with symmetry relaxation. This approach yielded reconstructions that, when low-pass filtered, contain obvious density that could be ascribed to GFP (**Figure 2B,C middle**). The fluorescent protein tags form an apparent cloud around the encapsulin core scaffolds, leading to particles with approximate diameters of 51 nm (50nm-GEMs) and 41 nm (40nm-GEMs) (**Figure 2B,C middle**). 2D projections of the previously determined *Quasibacillus thermotolerans* encapsulin map or a simulated map from the *Pyrococcus furiosus* crystal structure do not show this extra density (**Figure 2B,C right**) further suggesting this poorly resolved density can be attributed to the fused fluorescence tag. These maps demonstrate that endogenously expressed GEMs are primarily assembled as discrete particles with a particle diameter consistent with our expectations of 50 nm and 40 nm, respectively.

**Figure 2.**
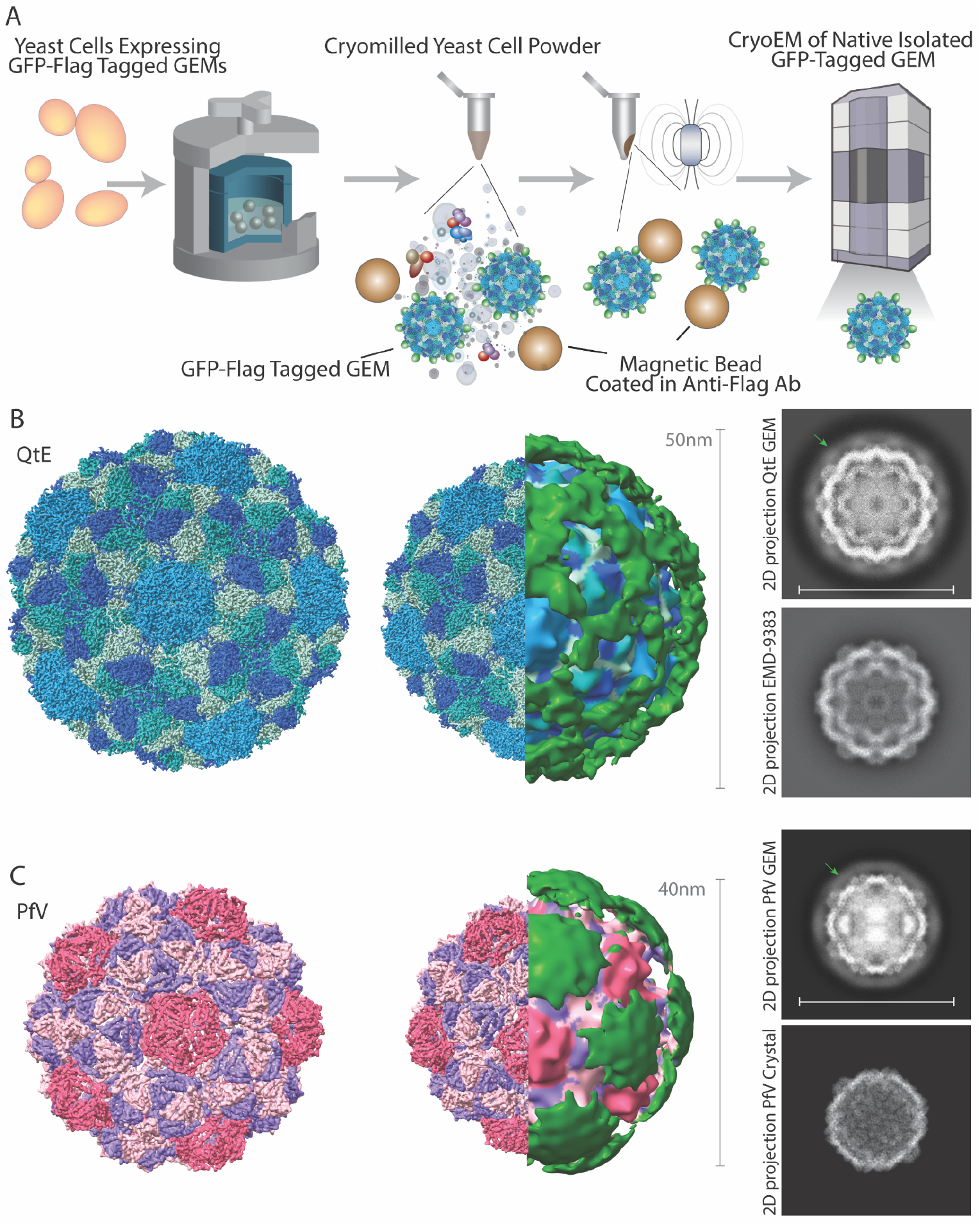
Isolation and structural characterization of GEMs from *S. cerevisiae*. **(A)** Schematic for isolation and structural analysis of *in vivo* assembled GEMs by cryo-EM SPA. *S. cerevisiae* expressing from the HIS3 promoter were grown, frozen and milled by a planetary ball mill. The resulting grindate was dissolved in an isolation buffer and GEMs were isolated using antibody-coated magnetic beads. GEMs eluted by FLAG peptide were then directly vitrified and imaged by cryo-TEM. **(B)** Cryo-EM density map of the *Quasibacillus thermotolerans* derived 50nm-GEM particle shown as a surface rendering (left) and as a cutaway view (middle). The low-pass filtered map (middle) reveals a labile outer layer, consisting of fluorescence proteins (green) enclosing a dense internal shell (blue). (right) 2D projections of the map from isolated 50nm-GEMs (top) show additional outer density (green arrow) not observed 2D projections in the previous SPA reconstruction of the *Quasibacillus thermotolerans* encapsulin scaffold (EMDB:9383, scale bar 50 nm). **(C)** Cryo-EM density map of the *Pyrococcus furiosus* derived 40nm-GEM particle shown as a surface rendering (left) and as a cutaway view (middle). The low-pass filtered map (middle) reveals a labile outer layer, consisting of fluorescence proteins (green) enclosing a dense internal shell (blue). (right) 2D projections of the map from isolated 40nm-GEMs (top) show additional outer density (green arrow) that is absent in 2D projections from a simulated map derived from the crystal structure of the observed 2D projections in the previous SPA reconstruction of the *Pyrococcus furiosus* iron storage scaffold (EMDB:9383, scale bar 40 nm).

**Supplemental Figure 2.**
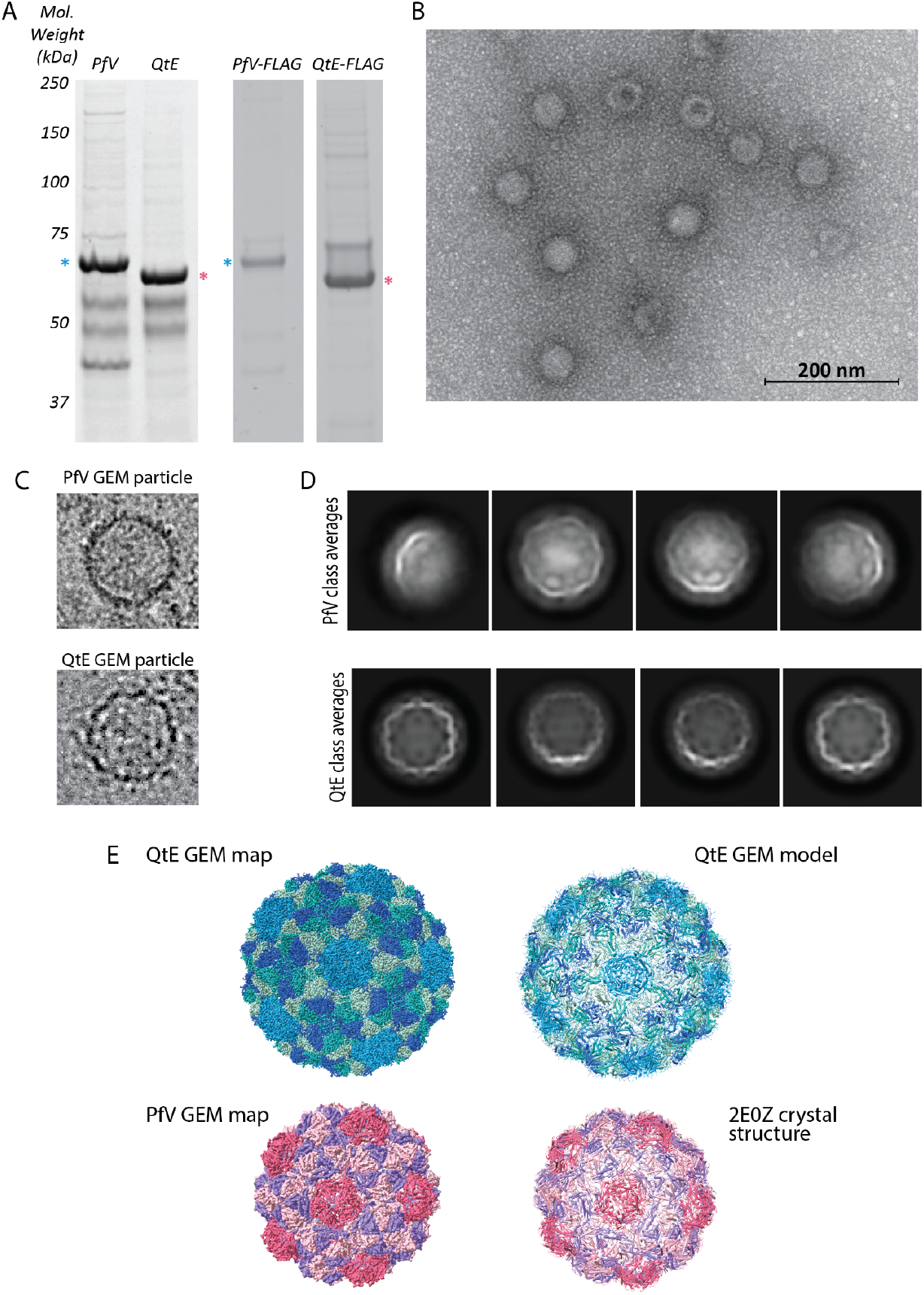
**(A)** SDS-PAGE analysis of affinity isolated 40nm-GEMs and 50nm-GEMs *in vivo* assembled from yeast. GEMs isolated by polyclonal llama GFP (left) or FLAG antibody (right) are quite pure and at the expected molecular weight. **(B)** Negative stain transmission electron microscopy of affinity isolated 50nm-GEMs showing homogenous particles of approximated expected size. **(C)** Example particles of the *Pyrococcus furiosus* (PfV) 40nm-GEM **(top)** and *Quasibacillus thermotolerans* (QtE) 50nm-GEM (**bottom**) taken from cryo-TEM micrograph. Particles are large and intact as expected. **(D)** 2D-class averages from PfV 40nm-GEMs **(top)** and QtE 50nm-GEMs (**bottom**) as calculated from cryoSPARC. Class averages show large intact GEMs, consistent with the particles in micrographs, with peripheral density clouds ascribable to a cloud of fluorescent protein tags. **(E)** Cryo-EM density maps and fit full-scale models of the QtE 50nm-GEMs **(top)** and PfV 40nm-GEMs (**bottom**).

**Supplemental Figure 3.**
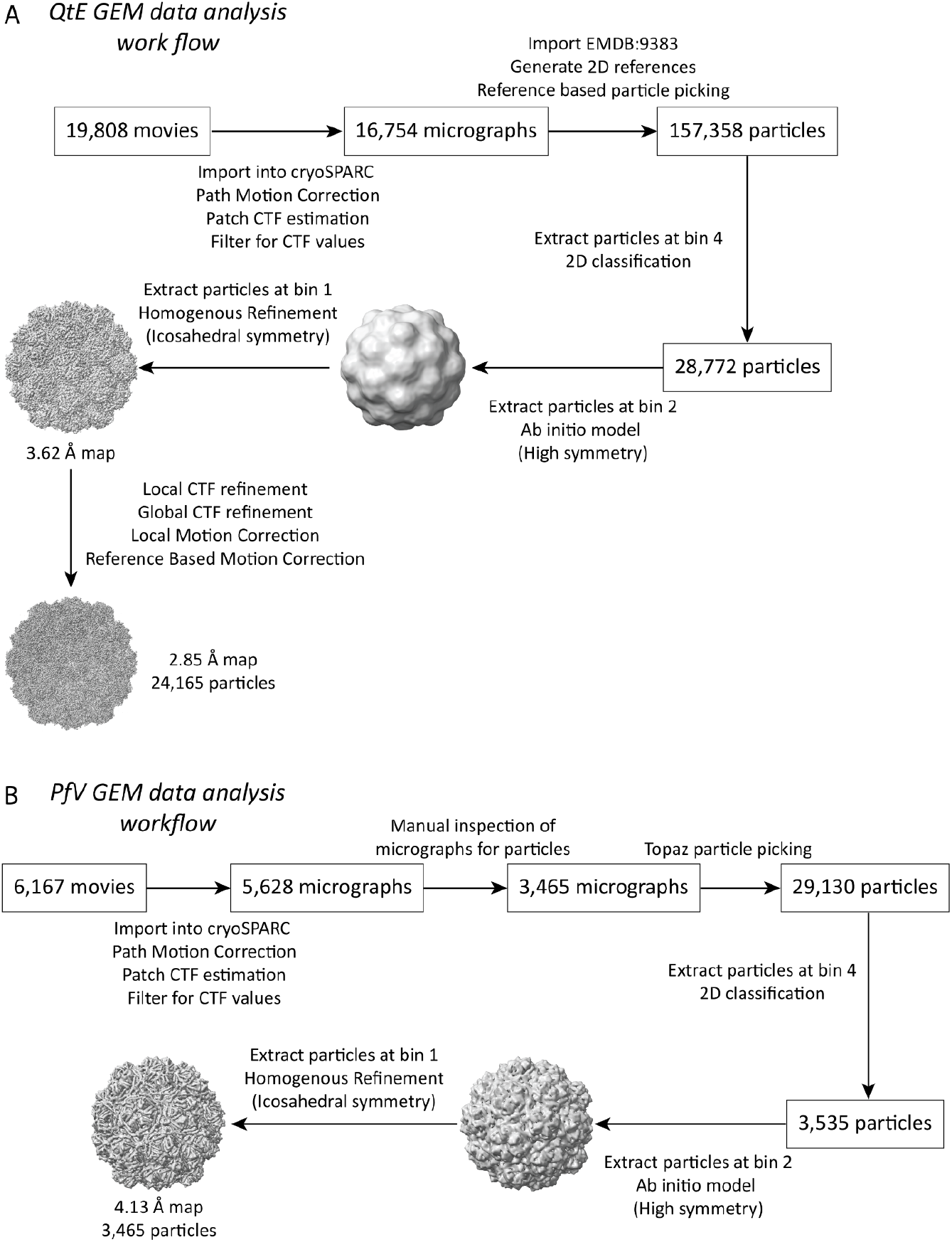
**(A)** Single particle analysis data processing workflow for QtE 50nm-GEM resulting in 2.85 angstrom reconstruction. **(B)** Single particle analysis data processing workflow for PfV 40nm-GEM resulting in 4.13 angstrom reconstruction.

### Structural Comparison of 40nm-GEMs and 50nm-GEMs

The 50nm-GEMs and 40nm-GEMs are formed by icosahedral assemblies of capsomeric proteins. The 50nm-GEM has a ~42 nm internal scaffold with icosahedral symmetry and T = 4 topology with 240 protomers per GEM. These protomers form 12 pentameric and 30 hexameric capsomers that occupy icosahedral vertices and faces (**Figure 3A**). The 2.85 angstrom map of the 50nm-GEM showed clear side-chain level features (**Supplementary Figure 4**). Using our improved higher-resolution map, we built an atomic model of the *in vivo* isolated *Quasibacillus thermotolerans* encapsulin protein. The four-protomer asymmetric unit from the endogenous GEM and the recombinant *Quasibacillus thermotolerans* encapsulin (PDB:6NJ8) are remarkably similar with an all atom root mean square deviation (RMSD) of 0.652 angstroms. The 40nm-GEM has a ~32 nm internal scaffold with icosahedral symmetry and T = 3 topology with 180 protomers per individual GEM assembly. These protomers form into 12 pentameric and 20 hexameric capsomers (**Figure 3B**) that occupy icosahedral vertices and faces. The limited resolution of this map precluded model building, however it exhibited an excellent fit with a previously determined crystal structure (PDB:2E0Z) and the biological assembly was consistent with the 3D reconstruction (**Supplementary Figure 2e)**.

**Figure 3.**
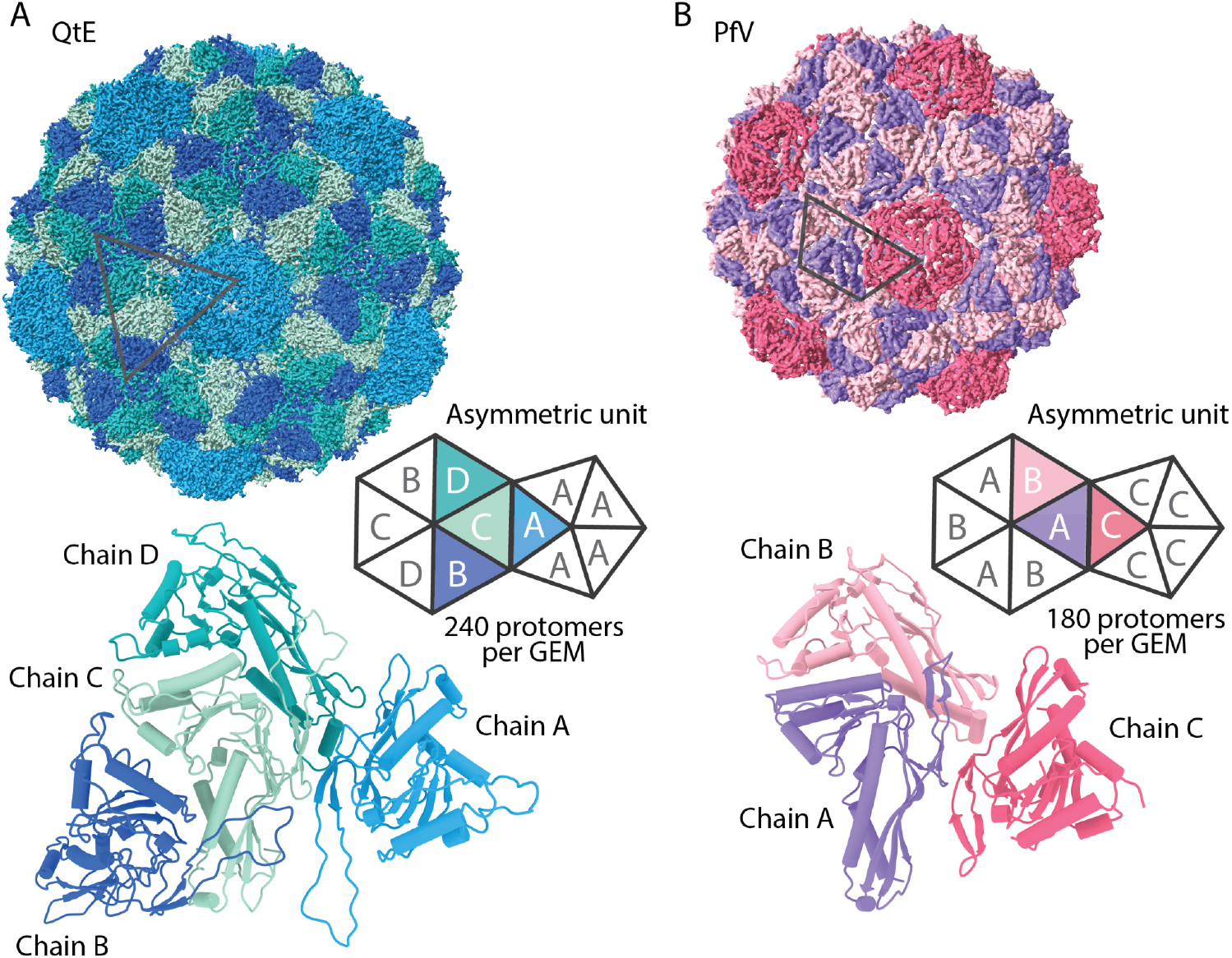
Structural arrangement of GEMs. **(A)** Cryo-EM map of the *Quasibacillus thermotolerans* encapsulin-derived 50nm-GEM pseudocolored by protomers. Four identical promoters (denoted chains A-D) of the encapsulin protein form an asymmetric unit. These asymmetric units form 12 pentameric and 30 hexameric capsomers that assemble in T = 4 icosahedral symmetry to form the GEM scaffold. **(B)** Cryo-EM map of the *Pyrococcus furiosus* encapsulin-derived 40nm-GEM scaffold pseudocolored by protomers. Three identical promoters (denoted chains A-C, fit from PDB: 2E0Z) form the asymmetric unit. These asymmetric units form 12 pentameric and 20 hexameric capsomers that assemble in T = 3 icosahedral symmetry to form the GEM scaffold.

**Supplemental Figure 4.**
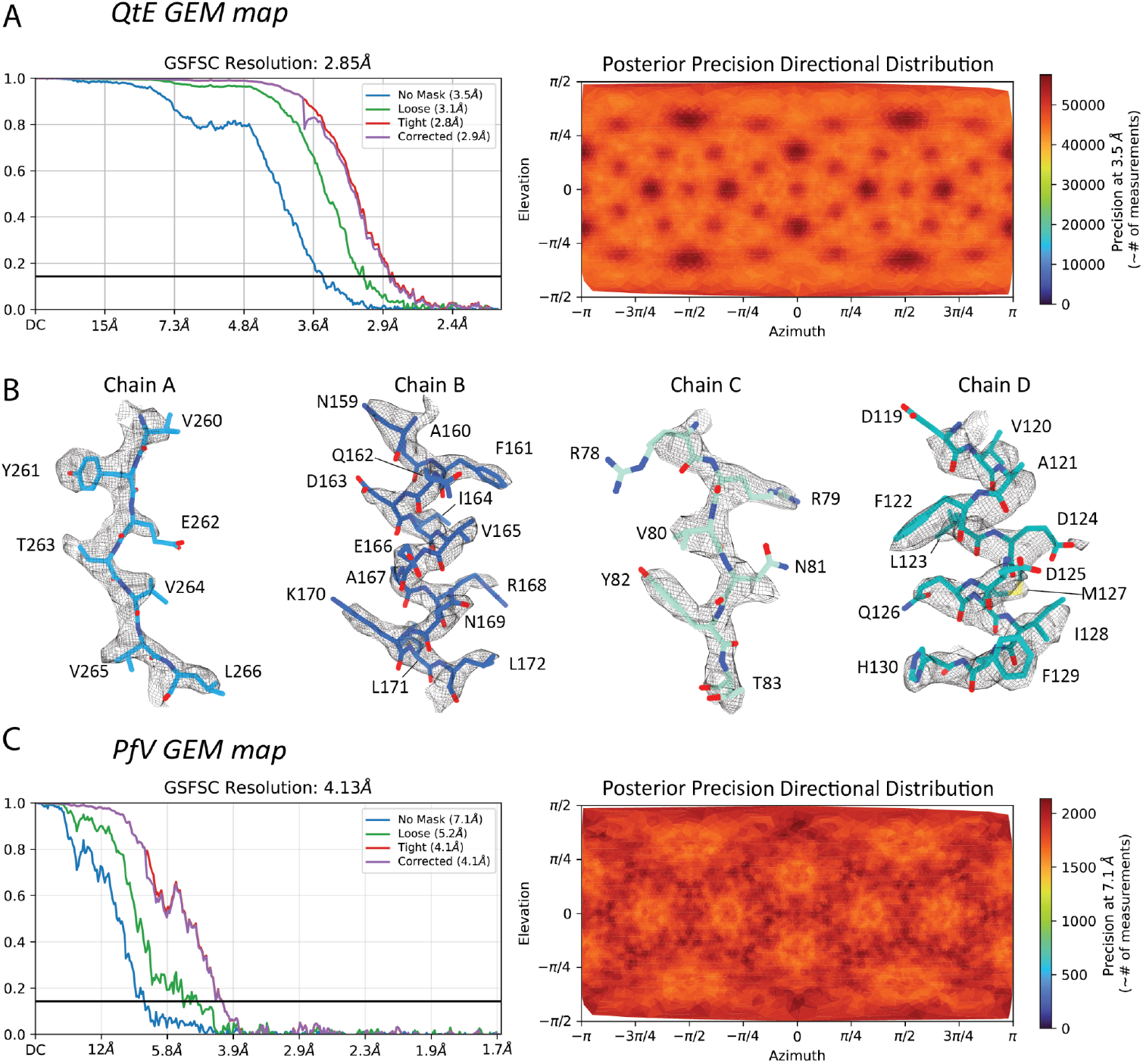
**(A)** Left: Fourier Shell Correlation (FSC) curve for the final 50nm-GEM reconstruction after icosahedral homogeneous refinement in cryoSPARC showing a final alignment resolution of 2.85 angstroms (FSC=0.143). Right: Posterior Precision Directional Distribution plot of the same refinement showing good angular coverage of the reconstruction. **(B)** Cryo-EM density with fit built models of the 50nm-GEMs shows clear side chain density of each chain. **(C)** Left: Fourier Shell Correlation (FSC) curve for the final 40nm-GEM reconstruction after icosahedral homogeneous refinement in cryoSPARC showing a final alignment resolution of 4.13 angstroms (FSC=0.143). Right: Posterior Precision Directional Distribution plot of the same refinement showing good angular coverage of the reconstruction.

### 50nm-GEMs probe a distinct diffusive regime compared to 40nm-GEMs in yeast cells

Following robust expression of GEMs in yeast, we quantified cytoplasmic nanoparticle diffusivity using live-cell imaging **(Supplementary Movies 1–2)** and single-particle tracking of 40nm-GEMs and 50 nm-GEMs. 40nm-GEM and 50nm-GEM expression plasmids were transformed to yeast cells and imaged **(Figure 4 A,B)**. Representative trajectories **(Figure 4 A,B)** demonstrate that both particle sizes diffuse throughout the cytoplasm. Yeast cells also maintain a particle density that is well suited for quantitative analysis. Single-particle trajectories revealed that 50nm-GEMs explored less cytoplasmic space than 40nm-GEMs. Notably, 50nm-GEMs diffuse more slowly than 40nm-GEMs, as indicated by diffusion-coded trajectories which show lower-diffusivity (blue) segments relative to the higher-diffusivity (red) segments characteristic of 40nm-GEMs.

**Figure 4.**
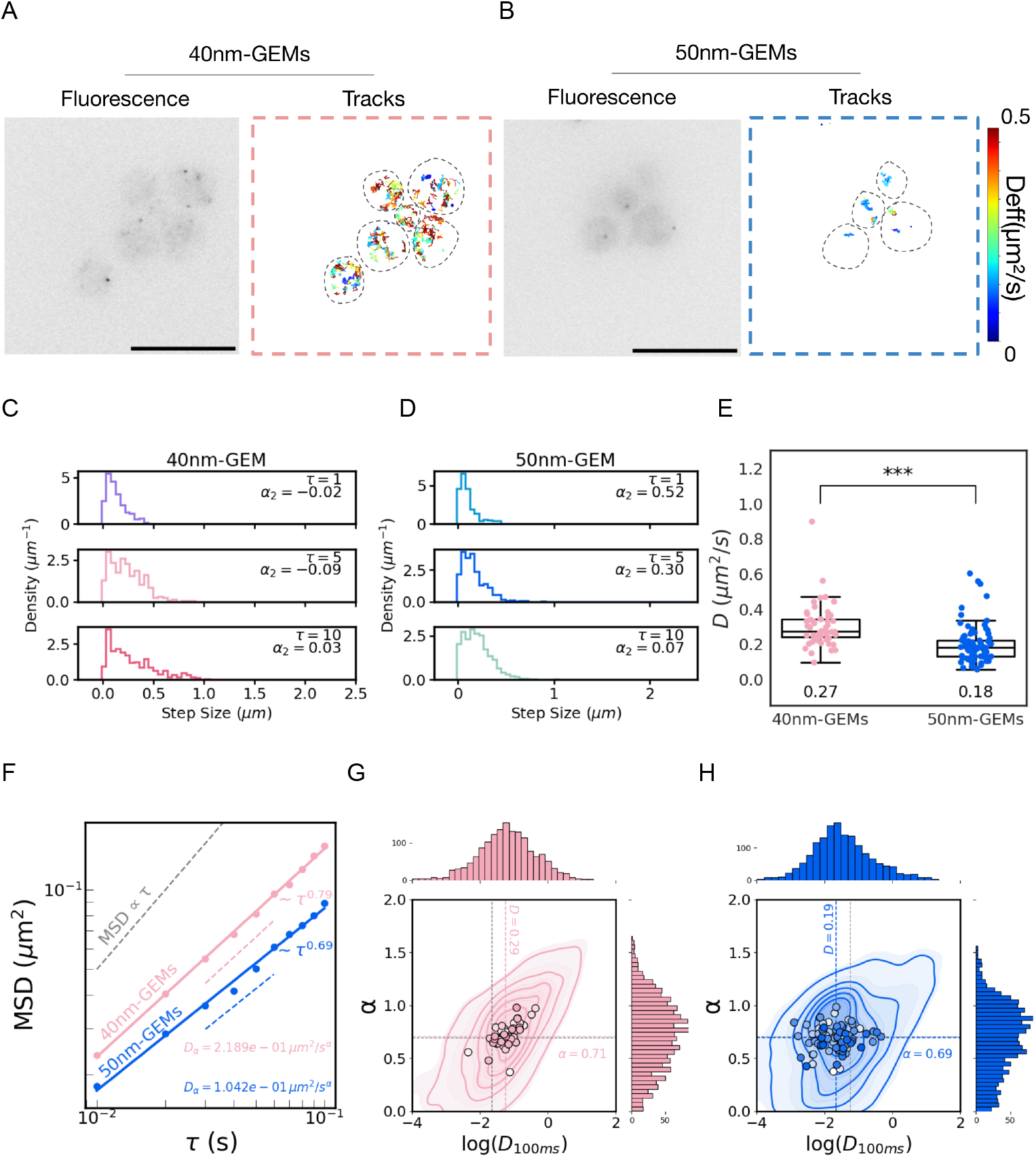
50nm-GEMs exhibit reduced diffusivity within the same diffusion mode in yeast. ***(S. cerevisiae)* (A–B)** Representative fluorescence images and single-particle tracking of 40nm-GEMs **(A)** and 50nm-GEMs **(B)**. Left panels show single-frame fluorescence images, illustrating the intracellular distribution of GEMs. Right panels show corresponding trajectories reconstructions, where individual tracks are color-coded by their effective diffusivity **D**. Warm colors indicate higher diffusivities, cool colors indicate lower diffusivities. Dashed outlines indicate segmented cellular regions used for tracking analysis. Scale bars are 10 microns. **(C–D)** Distributions of step sizes for 40nm-GEMs **(C)** and 50nm-GEMs **(D)** at increasing time lags (*τ*=1,5,10 frames). Each histogram is annotated with the corresponding non-gaussian parameter (α_2_), highlighting size- and time-dependent deviations from normal diffusion. **(E)** Quantification of effective diffusivities for 40nm-GEMs and 50nm-GEMs. Each dot represents an individual cell; box plots indicate median, interquartile range and 95% confidence intervals. Median **D** values are shown below each group. 40nm-GEMs exhibit significantly higher diffusivity than 50nm-GEMs (***, p<0.001). **(F)** Mean squared displacement (MSD) as a function of time lag*τ* for 40nm-GEMs (pink) and 50nm-GEMs (blue) shown on log–log axes. Dashed lines indicate power-law fits, MSD(*τ*)=4*D*_α_*τ*^α^, revealing subdiffusive behavior for both particle sizes. 40nm-GEMs exhibit both a higher diffusivity ***D***_α_and a larger anomalous exponent **α** compared to 50nm-GEMs. The gray dashed line indicates the scaling expected for normal diffusion ( MSD∝α). **(G–H)** Joint distributions of anomalous exponent (*α*) and effective diffusivity at (*τ* = 100) ms **D**_α_ for 40nm-GEMs **(G)** and 50nm-GEMs **(H)**. Individual points represent single cells; contour lines denote kernel density estimates of the trajectories. Histograms show the distributions of *log*(*D*_*α*_)(top) and *α* (right). Median values are indicated by dashed lines. 40nm-GEMs display a broader distribution shifted toward higher diffusivity and slightly larger *α*(median *α* ≈0.71) compared to 50nm-GEMs (median *α* ≈0.69). Yeast data include 24,594 tracked particles, with at least 300 cells per replicate analyzed across N = 3 independent biological replicates. Differences in median diffusivity were evaluated using a Mann–Whitney test (*p* < 0.001), whereas alpha values showed no statistically significant differences.

50nm-GEMs also exhibit a narrower and lower median step-size distribution than 40nm-GEMs across all time lags **(Figure 4C–D)**, consistent with the lower mobility of the larger particles. Gaussianity of the distribution was quantified with the α_2_ parameter (equation 1)

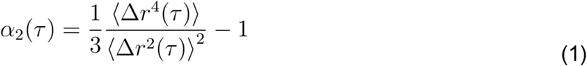

The α_2_ parameter is close to zero for both particle sizes, indicating approximately Gaussian step-size distributions.

Particle diffusivities were computed using the Einstein equation (equation 2) under the assumption of normal diffusion^32^,

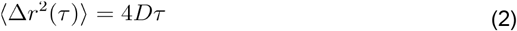

Under this framework, 50nm-GEMs exhibit markedly reduced mobility relative to 40nm-GEMs, with a significantly lower median effective diffusivity *D* for 50nm-GEMs ( *D*= 0.18 µm^2^ s^−^) compared to 40nm-GEMs (*D* = 0.27 µm^2^ s^−^; p < 0.001; **Figure 4E**).^32,33^

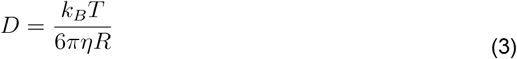

We next quantified the anomalous diffusion exponent α to determine whether GEM dynamics deviate from normal diffusion by fitting the the time- and ensemble-averaged mean-squared displacement (MSD) to the anomalous diffusion model:

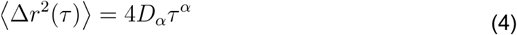

In log-log scale Diffusivity (D) is proportional to the y-intercept and the anomalous diffusion exponent (α) is proportional to the slope of the fit. The anomalous diffusion exponent α quantifies these deviations, with α=1 corresponding to normal diffusion, α<1 to subdiffusive motion arising from constraints such as crowding, transient binding, or confinement, and α>1 to superdiffusive behavior typically associated with persistent or active transport. Thus, α captures information about the physical environment that is not contained in the diffusion coefficient alone.

Both 40nm-GEMs and 50nm-GEMs displayed apparently subdiffusive dynamics, with fitted anomalous exponents (α = 0.71) and (α = 0.69), respectively **(Figure 4F)**. However, 3D simulations incorporating finite focal depth and 2D projection demonstrate that similar apparently subdiffusive exponents can arise from purely Brownian motion due to the limited focal plane **(Supplementary Figure 1C)**. Indeed, simulations of Brownian motion using the measured diffusivities of GEMs gives α ~ 0.8. Therefore, GEMs have only slightly subdiffusive motion. The close agreement in α values between the two probe sizes also suggests that both GEMs experience comparable mild interactions with their local environment

To quantify particle- and cell-to-cell heterogeneity, we next fit the parameters of the anomalous diffusion equation for each individual trajectory. Because the number of trajectories is large, we visualized the resulting parameter space using Kernel Density Estimates (KDEs) **(Fig. 4G–H)**, and median values per cell are additionally indicated by circles. These plots revealed higher cell-to-cell and intracellular heterogeneity of diffusivities for 50nm-GEMs compared to 40nm-GEMs. Notably, the anomalous exponent α was comparable between particle sizes. The distributions of track-level data were unimodal rather than bimodal (see histograms above and to the right of scatter-plots), arguing against the presence of distinct dynamical subpopulations.

### Engineering 50nm-GEMs for assembly in mammalian cells

After demonstrating the ability to express 50nm-GEMs in *S. cerevisiae*, we expanded this tool for use in mammalian cells. We subcloned the 50nm-GEM cassette from the yeast expression plasmid into a lentiviral backbone (from plasmid pLH1984-Holt lab database-a pLVX-based pUBC-PfV-6xGS-Sapphire construct), thereby replacing the original 40nm-GEM while preserving the pLVX-backbone and UBC promoter. Upon imaging with confocal microscopy (see methods), we did not see 50nm-GEM assembly, but instead observed dispersed green fluorescence throughout the cells, and small, rapidly diffusing subassemblies. We hypothesized that this failure to assemble was due to the presence of a nucleation barrier that hindered effective 50nm-GEM particle formation in the mammalian cytoplasm. Attempts to overcome this barrier by increasing DNA concentration or enhancing crowding and monomer concentration through sorbitol treatment were unsuccessful.

We reasoned that more efficient nucleation could be achieved by locally concentrating monomers via an additional interaction. To test this idea, we fused a short peptide from the *Caulobacter crescentus* PopZ protein^34^ to the N-terminus of the 50nm-GEM coding sequence via a GS linker. Specifically, we used the H3 C-terminal helical oligomerization domain, which has previously been described to be sufficient for condensate formation^35^. Transient transfection of HeLa cells with this construct resulted in efficient assembly of 50nm-GEM nanoparticles in the cytoplasm (**Figure 5**).

**Figure 5.**
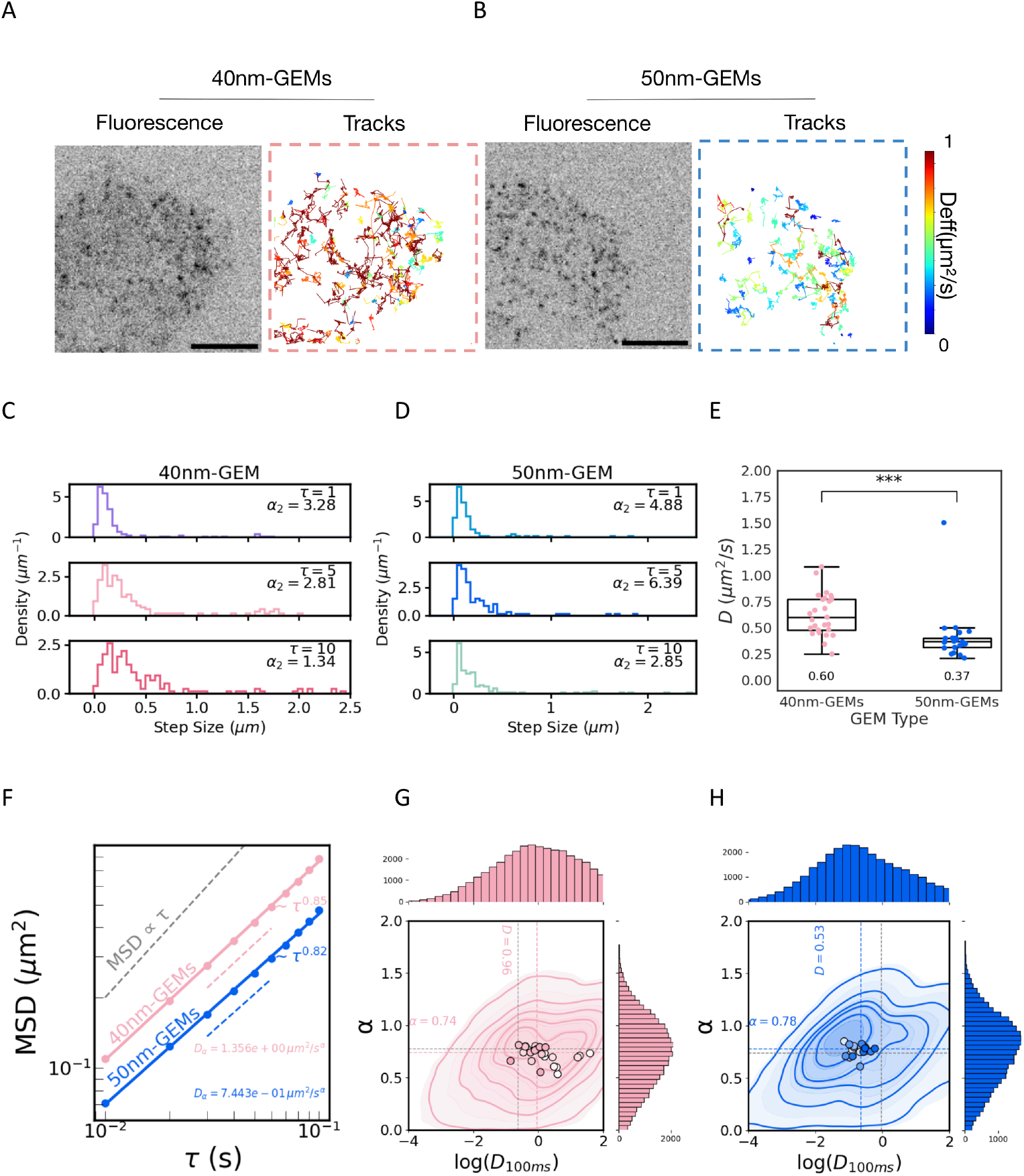
50nm-GEMs exhibit reduced diffusivity within the same diffusion mode in mammalian (HeLa) cells. Representative fluorescence images and single-particle tracking of 40nm-GEMs (**A**) and 50nm-GEMs (**B**). Left panels show single-frame fluorescence images, illustrating the intracellular distribution of GEMs. Right panels show corresponding trajectories reconstructions, where individual tracks are color-coded by their effective diffusivity *D*. Warm colors indicate higher diffusivities, cool colors indicate lower diffusivities. Dashed outlines indicate segmented cellular regions used for tracking analysis. Scale bars are 10 microns. (Number of tracks filtered to show a similar number of tracks on both panels) **(C–D)** Distributions of step sizes for 40nm-GEMs (**C**) and 50nm-GEMs (**D**) at increasing time lags (*τ* =1,5,10 frames). Each histogram is annotated with the corresponding non-Gaussian parameter (α_2_), highlighting size- and time-dependent deviations from normal diffusion. **(E)** Quantification of effective diffusivities for 40nm-GEMs and 50nm-GEMs. Each dot represents an individual cell; box plots indicate median, interquartile range and 95% confidence intervals. Median *D* values are shown below each group. 40nm-GEMs exhibit significantly higher diffusivity than 50nm-GEMs (***, p<0.001). (**F**) Mean squared displacement (MSD) as a function of time lag *τ* for 40nm-GEMs (pink) and 50nm-GEMs (blue) shown on log–log axes. Dashed lines indicate power-law fits, MSD(*τ*)=4*D*_*α*_*τ* ^*α*^ revealing subdiffusive behavior for both particle sizes. 40nm-GEMs exhibit both a higher effective diffusion coefficient (D) and a larger anomalous exponent *α* compared to 50nm-GEMs. The gray dashed line indicates the scaling expected for normal diffusion (MSD ∝ *τ*). **(G–H)** Joint distributions of anomalous exponent *α* and effective diffusivity at *τ* =100 ms for *D* 40nm-GEMs (**G**) and 50nm-GEMs (**H**). Individual points represent single cells; contour lines denote kernel density estimates of the trajectories. Histograms show the distributions of *log* (*D*_*α*_)(top) and (right). Median values are indicated by dashed lines. 40nm-GEMs display a broader distribution shifted toward higher diffusivity and slightly larger (median α ≈ 0.85) compared to 50nm-GEMs*α* (median α ≈ 0.82). Mammalian data from 91,333 tracks, from ≥50 cells per replicate across N = 3 biological replicates. Statistical significance of median diffusivity differences was assessed using a Mann–Whitney test (*p* < 0.001). Alpha values were not significantly different.

### 50nm-GEMs exhibit slower diffusion than 40nm-GEMs in mammalian cells

Following successful expression of GEMs in HeLa cells, we performed single-particle tracking of 40nm-GEMs and 50nm-GEMs **(Supplementary Movies 3-4)**. 50nm-GEM and 40nm-GEM expression plasmids were transiently expressed in HeLa cells **(Figure 5A,B)** and imaged 24 hours after transfection. Compared to yeast, mammalian cells contained a substantially higher number of particles **(Figure 5A,B and Fig. 4A,B)**.

Because increased particle density can lead to tracking errors, particularly due to particle misidentification, we performed simulations to quantify errors arising from high-density conditions **(Supplementary Fig. 1)**. These simulations defined a particle-density threshold **(Supplementary Fig. 1B bottom)** below which tracking remained reliable; consequently, all subsequent analyses were restricted to cells with trajectories within this density range.

Both particle sizes diffuse throughout the cytoplasm but trajectories revealed that 50nm-GEMs explored less space than 40nm-GEMs, consistent with observations in yeast **(Figure 5A,B)**. 50nm-GEM trajectories tended to have lower diffusivity (blue) than the higher-diffusivity (red) tracks characteristic of 40nm-GEMs, indicating a size-dependent slowdown of particle motion.

Step-size distributions were narrower, with lower median displacements for 50nm-GEMs across all time lags **(Figure 5 C,D)**, consistent with the expected reduction in mobility for larger particles. The non-Gaussian parameter α_2_ **(Figure 5 C-D)** indicated moderate deviations from Gaussian behavior (moreso than yeast **Figure 4 C-D**) at all time lags, suggesting that motion is somewhat heterogeneous.

Time- and ensemble-averaged mean squared displacements (MSDs) **(Figure 5F)** showed substantially higher diffusivities in mammalian cells than in yeast: D was 1.36 and 0.74 µm^2^ s^-1^ for 40nm-GEMs and 50nm-GEMs, respectively.

Again both 40nm-GEMs and 50nm-GEMs were apparently subdiffusive with α = 0.85, α = 0.82, respectively **(Figure 5F)**. However, these values are almost the same as the α ~ 0.8 that arises simply from the artifact of particles being confined within a limited focal plane **(Supplementary Figure 1C)**.

We next quantified individual trajectory parameters, including effective diffusivity (D) and anomalous exponent (α), and visualized them in two-dimensional space. Kernel density estimation (KDE) was used to show distributions of individual track behavior, and median values per cell are indicated by circles **(Figure 5 H,I)**. Tracks showed a broader distribution of diffusivities compared to yeast. Compared to yeast, α values were slightly higher (α ≈ 0.8 for both particles). Therefore, the mammalian cytoplasm exhibits more heterogeneity than yeast while remaining largely diffusive.

Fluorescence intensity distributions were influenced by particle number: low particle numbers produced sparse, broad signals, while high densities yielded smoother, approximately Gaussian distributions due to overlapping signals and averaging effects **(Supplementary Fig. 1B, center)**.

Because we had established a simulation framework, we sought to explore how optical artifacts could influence the measured fluorescence intensity within cells. Properly assembled nanoparticles are expected to exhibit a narrow, unimodal fluorescence intensity distribution reflecting uniform stoichiometry; we examined the brightness distribution of tracked particles across cell types. In yeast, intensity distributions were tightly unimodal **(Fig. 1B left)**, consistent with homogeneous particle assembly. In contrast, mammalian cells displayed broader distributions **(Figure 1B right)**: 40 nm GEMs were moderately broadened relative to yeast, whereas 50 nm GEMs exhibited similarly broad profiles with additional peaks, suggesting potential heterogeneity in the detected signal.

Given that the nanoparticles are structurally identical across systems, we reasoned that differences in imaging conditions—particularly high particle density leading to overlapping particles—could underlie this effect. To test this possibility, we used simulated data **(Supplementary Fig. 1B)** in which particle density was systematically varied to quantify its impact on the resulting intensity distributions. The simulations revealed that low densities **(Supplementary Figure 1B, center left)** produced a single peak of intensity estimates and as particle density increased **(Supplementary Figure 1B, mid-center)** a second peak emerged due to detections of two overlapping particles. Very high densities led to brighter, multimodal intensity profiles due to multiple particles overlapping **(Supplementary Figure 1B, center right)**. Together, these results indicate that the multimodality observed in mammalian cells can arise from density-dependent imaging artifacts rather than intrinsic defects in nanoparticle assembly.

Together, these results indicate that 50nm-GEMs diffuse more slowly than 40nm-GEMs in the mammalian cytoplasm. Compared to yeast, the mammalian cytoplasm shows stronger non-Gaussian behavior and slightly greater heterogeneity, suggesting that the larger and more complex cytoplasmic architecture of mammalian cells imposes spatially varying constraints on particle motion.

## Discussion

In this study we developed, structurally characterized, and functionally validated **50nm-GEMs** as a new tool for intracellular nanorheology. By using encapsulin-based scaffolds from *Quasibacillus thermotolerans*, we generated a nanoparticle that self-assembles *in vivo*, is about two times brighter than its 40 nm predecessor, and probes a larger length scale that is highly relevant to mesoscale cell biology. Our work demonstrates that these probes assemble with high fidelity in *S. cerevisiae* and, after additional engineering, also in mammalian cells, enabling systematic cross-species comparisons of intracellular physical properties at the 50 nm length-scale.

### Preservation of large macromolecular structures in single step affinity isolation

Methods for single step affinity isolation of macromolecular assemblies for cryo-EM SPA have typically leveraged antibody immobilized grids^36,37^or particle electrostatics^38,39^. Our own efforts in this vein have focused primarily on the Nuclear Pore Complex (NPC)^28^, a 50 MDa multimeric complex that is significantly larger than most macromolecular complexes. Here we show this single step affinity isolation of complexes from cryomilled cells is amenable to a broader set of structural targets on a size scale of common macromolecular machines. This technique preserves near-atomic structural fidelity of isolated complexes for high resolution structural determination. Structures of the *in vivo* assembled 40nm-GEMs and 50nm-GEMs confirmed that they assemble into the expected T=3 and T=4 icosahedral symmetries. This confirms that orthologous large protein assemblies can be transferred across species and faithfully reconstituted *in vivo*, validating the portability of GEM designs, and confirming their preservation as intact, native architectures during single-step affinity isolation.^28,3836–40^

### A robust, bright mesoscale probe derived from a native T = 4 encapsulin

Cryo-EM reconstruction of the *in vivo* assembled 50nm-GEMs confirmed that the encapsulin scaffold assembles into a ~42 nm protein shell that is surrounded by a flexible “cloud” of GFP proteins adding ~8–10 nm to the overall diameter — consistent with a ~50 nm hydrodynamic radius. Importantly, the GFP density was only recoverable when symmetry constraints were relaxed, indicating substantial structural mobility of the peripherally tethered fluorophores. This flexibility likely improves probe isotropy during diffusion, while the high monomer count per GEM underlies the two-fold brightness increase relative to 40nm-GEMs. This enhanced brightness directly addresses a key challenge in mesoscale nanorheology: the requirement for highly sensitive microscopes capable of acquiring high-frame-rate data. The significantly improved signal-to-noise ratio of 50nm-GEMs makes them compatible with more standard imaging platforms, enabling broader adoption of live-cell nanorheology.

### Size-dependent cytoplasmic diffusion across species

Single-particle tracking in both *S. cerevisiae* and HeLa cells revealed the expected size-dependent decrease in diffusivity, with 50nm-GEMs consistently moving more slowly and exhibiting reduced step sizes relative to 40nm-GEMs. Crucially, despite their reduced mobility, 50nm-GEMs exhibited similar anomalous exponents (*α*) relative to 40nm-GEMs, indicating that the mode of cytoplasmic diffusion remains unchanged across probe sizes. In both cases this motion is characterized by slightly subdiffusive, viscoelastic motion. Thus, GEM size primarily tunes the effective viscosity experienced by the particle, rather than altering its qualitative interaction with the intracellular material environment.

### 50nm-GEMs as a new tool to interrogate mesoscale cytoplasmic organization

40nm-GEMs have already proven to be a powerful tool for dissecting how growth conditions, nutrient signaling, and aging^12,41,42^ alter cytoplasmic crowding, metabolic activity, and mesoscale organization^10,43,44^. The introduction of 50nm-GEMs now extends this capability to a larger, biologically crucial length scale. Because many key molecular assemblies fall within the ~40–100 nm range, 50nm-GEMs may be especially sensitive to changes in condensate abundance, organelle surface properties, polysome organization, or cytoskeletal mesh size. Larger GEMs may also better detect spatial heterogeneity in the cytoplasm—something suggested by the broader diffusion parameter distributions we observe.

Finally, the higher brightness of 50 nm GEMs enables tracking at 30–100 Hz using lower laser powers and potentially standard widefield and epifluorescence microscopy, reducing the need for highly specialized imaging platforms. This increased accessibility facilitates high-throughput intracellular nanorheology, enabling genetic and chemical screens for regulators of cytoplasmic mechanics, longitudinal stress-response measurements, and comparative biophysical studies across model organisms that were previously difficult to perform at scale.

### Conclusions and outlook

We report the design, structural elucidation, and functional characterization of a genetically encoded ~50 nm intracellular nanoparticle optimized for mesoscale nanorheology. Compared to previous generations of GEMs, these probes are substantially brighter, enabling robust and reproducible tracking. They are compatible with expression in both yeast and mammalian cells and are sensitive to biophysical features at a mesoscale length range occupied by many key intracellular assemblies. Importantly, their increased brightness and size yields longer particle trajectories, enabling analysis over extended time windows and allowing more reliable extraction of intrinsic physical properties of the intracellular environment.

Together, our results establish 50nm-GEMs as a versatile new tool for probing the physical organization of living cells. Looking ahead, combining GEMs of multiple sizes, colors, and compartments with advanced analysis frameworks will allow researchers to map the biophysical landscape of the cell interior with unprecedented resolution to reveal how cells regulate their internal physical state in health, differentiation, stress, and disease.

## Methods

### Yeast strains

All *S. cerevisiae* strains constructed for and used in this study are listed in Table 1. Unless otherwise stated, strains were grown at 30 °C in synthetic complete media + 2% dextrose (SCD) according to standard Cold Spring Harbor Protocols^45^. Yeast strains were constructed using standard molecular genetic methods, and verified by Oxford Nanopore sequencing and fluorescence microscopy for GEM expression. In preparation for imaging via fluorescence microscopy, cells were grown at 30°C in a rotating incubator overnight. The next morning, cells were grown for 4-5 hours to reach log phase, at which point they reached an OD600 between 0.2 and 0.4.

**Table 1.**
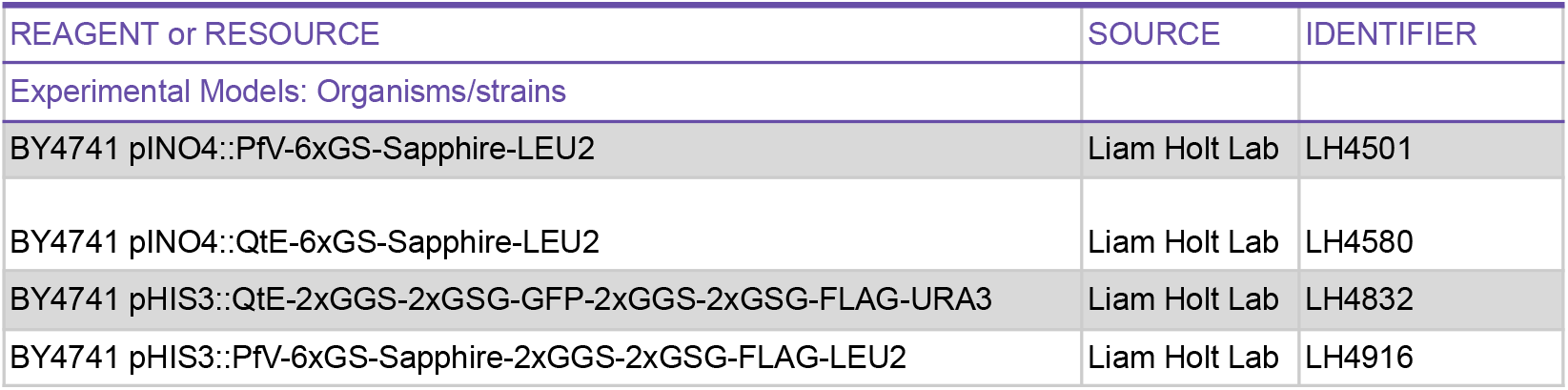
Yeast Strain Information.

**Table 2.**
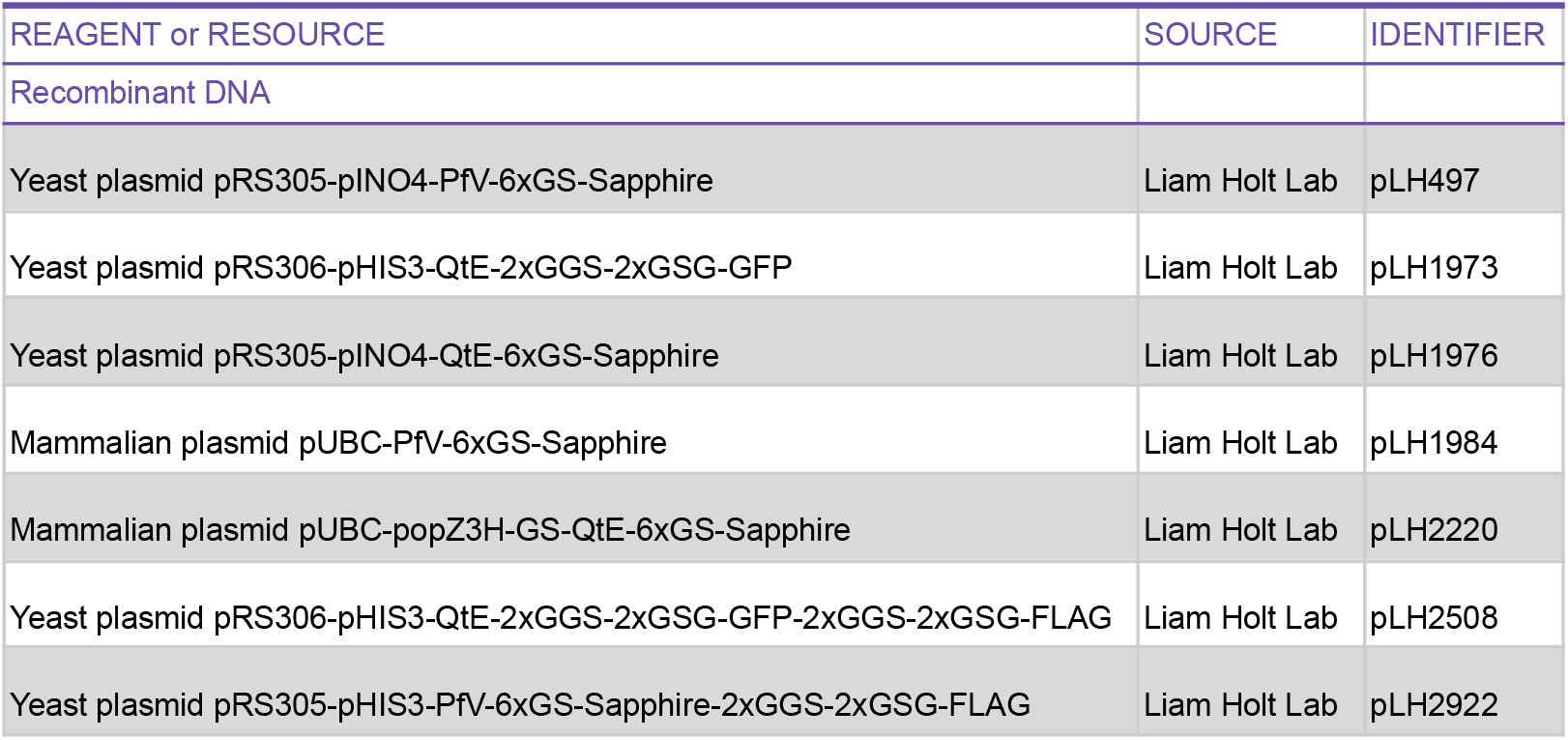
Plasmid information.

| REAGENT or RESOURCE | SOURCE | IDENTIFIER |
| --- | --- | --- |
| Recombinant DNA |  |  |
| Yeast plasmid pRS305-pINO4-PfV-6xGS-Sapphire | Liam Holt Lab | pLH497 |
| Yeast plasmid pRS306-pHIS3-QtE-2xGGS-2xGSG-GFP | Liam Holt Lab | pLH1973 |
| Yeast plasmid pRS305-pINO4-QtE-6xGS-Sapphire | Liam Holt Lab | pLH1976 |
| Mammalian plasmid pUBC-PfV-6xGS-Sapphire | Liam Holt Lab | pLH1984 |
| Mammalian plasmid pUBC-popZ3H-GS-QtE-6xGS-Sapphire | Liam Holt Lab | pLH2220 |
| Yeast plasmid pRS306-pHIS3-QtE-2xGGS-2xGSG-GFP-2xGGS-2xGSG-FLAG | Liam Holt Lab | pLH2508 |
| Yeast plasmid pRS305-pHIS3-PfV-6xGS-Sapphire-2xGGS-2xGSG-FLAG | Liam Holt Lab | pLH2922 |

### Construction of 50nm-GEM plasmids

The open reading frame (ORF) encoding the *Quasibacillus thermotolerans* (QtE) encapsulin protein, based on its published cryo-EM model ([PDB:6NJ8](https://www.rcsb.org/structure/6NJ8)), was codon-optimized for yeast expression and synthesized as an IDT gene block ([IDT](http://www.idtdna.com/pages)). We utilized PCR and Gibson assembly to replace the 40nm-GEM ORF from the yeast expression vector pLH497 with the gene block of the 50nm-GEM ORF. The resulting plasmid was pLH1976:pRS305-pINO4-*QtE*-GS-Sapphire.

#### GEM-Sapphire construct

The fluorescent nanoparticle used in this study incorporates a variant of the Sapphire variant of green fluorescent protein (FPbase ID: G4VP2^25^). Sequence alignment revealed two amino acid substitutions relative to the canonical Sapphire protein: H231→T and M233→I (numbering based on the reference sequence). Residue H231 matches the corresponding residue in wild-type GFP and has not been associated with altered fluorescence properties. Residue M233 is solvent-exposed and outside the hydrophobic core; the M→I substitution is conservative, as both residues are nonpolar and hydrophobic. Based on structural considerations, these substitutions are unlikely to substantially affect fluorophore behavior.

To facilitate GEM purification for cryo-EM analysis we used Gibson Assembly to engineer a yeast expression plasmid under the HIS3 promoter, thereby increasing the number of particles expressed per cell. This was done by whole plasmid amplification of pLH1973:pRS306-pHIS3-QtE-GS-GFP using primers with tails containing the FLAG sequence, followed by T4 PNK Phosphorylation and T4 ligation. The resulting plasmid was pLH2508:pRS306-pHIS3-QtE-GS-GFP-GS-FLAG.

For mammalian expression, the *QtE*-GS-*T*-Sapphire gene cassette from vector pLH1976 was subcloned into a *pLVX* lentiviral backbone under the control of the *pUBC* promoter. Initial versions of the 50nm-GEMs did not assemble efficiently. To increase assembly efficiency, we used Gibson Assembly to add the H3 oligomerization domain of the PopZ protein from *Caulobacter crescentus* to the N-terminus of the 50nm-GEM ORF via a GS linker.

All plasmids were stored as bacterial glycerol stocks, and those generated in this study are available through AddGene (Accession No. XXXX).

#### Mammalian cell culture and transfection

HeLa cells were cultured in high-glucose Dulbecco’s Modified Eagle Medium (DMEM; Gibco) supplemented with 10% fetal bovine serum (FBS; Gemini Bio), penicillin (100 U/ml) and streptomycin (100 µg mL(^{-1})). Cells were maintained at 37°C in a humidified incubator with 5% CO_2_. For imaging experiments, cells were plated on 6-well glass-bottom dishes (Cellvis) at a density of 1 × 10(^5) cells per well and grown to 60–80% confluency within 24 h. Cells were transiently transfected using FuGENE® HD Transfection Reagent (Promega) according to the manufacturer’s protocol. Briefly, 0.5 µg of transfection-grade plasmid DNA (purified using the ZymoPURE™ Plasmid Miniprep Kit; Zymo) was mixed with 1 µL FuGENE HD in 200 µL Opti-MEM Reduced Serum Medium (Gibco). The transfection mixture was incubated at room temperature for 10 min and then added dropwise to cells. After 24 h, the culture medium was replaced with fresh DMEM, and cells were imaged for GEM expression. A plasmid concentration of 0.5 µg per well yielded optimal GEM expression and was used for all experiments. HeLa cells used in this study were kindly provided by Prof. Jef Boeke (Institute for Systems Genetics, NYU Langone).

#### Affinity-purification of GEMs from *S. cerevisiae*

We adapted a previously published method for the isolation of endogenous whole NPCs from *S. cerevisiae*^*46*^ to purification of natively expressed GEMs. Briefly, strains harboring FLAG-tagged 40nm-GEMs and 50nm-GEMs incorporated at the *LEU2* locus were grown in YPD media at 30°C until mid-log phase (~3×10^7^ cells/ml), harvested, frozen in liquid nitrogen and cryogenically lysed in a planetary ball mill PM 100 (Retsch) (http://lab.rockefeller.edu/rout/protocols). Affinity purification was performed using anti-FLAG or anti-GFP dynabeads (Thermo) in clarified lysate of cryo-milled yeast in resuspension buffer (20 mM HEPES/KOH pH 7.4, 500 mM sodium chloride, 0.1% (w/v) Triton X-100, 1 mM DTT, 10% (v/v) glycerol, 1/500 (v/v) Protease Inhibitor Cocktail (Sigma)). GEMs were either eluted by denaturing for SDS-PAGE analysis or FLAG peptide for structure characterization. For SDS-PAGE analysis, affinity captured GEMs were eluted from beads by addition of 20 μl of 1x LDS (lithium dodecyl sulfate) loading buffer (Thermo Fisher) and vortexing for 10 minutes at room temperature. Eluted GEMs were run on an NuPAGE 4-12% gel for 60 minutes and analyzed by Coomaisse. ^46^

For structural studies, GEMs were eluted by nutating beads with elution buffer (20 mM HEPES/KOH pH 7.4, 500 mM sodium chloride, 0.1% (w/v) Tween-20,1 mM DTT) supplemented with 1mg/mL of 3xFLAG peptide for 20 minutes at room temperature. For 50nm-GEMs Eluted GEMs were further analyzed by gradient sedimentation through a 10-60% glycerol gradient with a SW60 Ti (Beckman) rotor. Gradients were spun at 32,000 RPM for 2 hours and 45 minutes and fractionated and profiled using BioComp piston fractionator and Triax profiler. Peak fractions, as determined by fluorescence signal, were pooled and concentrated.

#### Cryo-EM analysis of GEMs

Grids for single particle analysis were made by incubating R2/2 300 mesh Quantifoil grids with thin carbon support on 20 µL drops from concentrated GEMs for 10 minutes in a humidity chamber to allow for particle adhesion. Grids were then washed with wash buffer (20 mM HEPES/KOH pH 7.4, 150 mM sodium chloride, 1 mM DTT) ensuring to keep the back of the grid dry. A final volume of 3 µL of wash buffer was applied to grids in the Vitrobot (ThermoFisher) humidity chamber. Grids were blotted for 3 seconds at blot force 3 and plunged using a Vitrobot.

For SPA analysis of 50nm-GEM grids were imaged on a Titan Krios (ThermoFisher) with spherical aberration (Cs) correction and BioQuantum energy filter (Gatan), using a K3 direct electron detector operated in super-resolution mode. Movies of 40 frames were collected at a nominal pixel size of 0.54 angstroms with a target electron flux of 51 e^-^/Å^2^. Movies were imported into cryoSPARC v4.4.1 ^47^ and binned by 2 during patch motion correction for a final pixel size of 1.08 angstroms. Motion corrected micrographs were CTF corrected with patch CTF estimation and were manually filtered for quality of CTF fit, total micrograph motion and ice quality, yielding a dataset of 7,323 micrographs. For SPA analysis of 40nm-GEMs, grids were imaged on EF-Krios operated at 300 kV with a GatanK3 imaging system collected at 105,000X nominal magnification. The calibrated pixel size of 0.4130 Å was used for processing. Movies were collected using Leginon (Suloway et al., 2005) at a dose rate of 30.82 e^-^/Å^2^/s with a total exposure of 1.80 seconds, for an accumulated dose of 55.48 e^-^/Å^2^. Intermediate frames were recorded every 0.04 seconds for a total of 45 frames per micrograph. A total of 6167 images were collected at a nominal defocus range of −0.1 – 2.8 μm. Motion correction and CTF estimation was performed as described above, including binning during motion correction for a working pixel size of 0.826 angstroms per pixel.

Due to their significant size, both 40nm-GEMs and 50nm-GEMs were easily distinguished on micrographs. For 50nm-GEMs, 2D references were generated from the previously reported cryo-EM map (EMDB:9383) and picked by reference based picking in cryoSPARC, yielding 157,358 particles that were subject to 2D classification. Even in initial 2D classes, fuzzy density outside of the clearly detailed core scaffold was observed, suggesting C-terminally linked GFPs were oriented on the external surface and provided extra volume to the 50nm-GEM. Only 2D classes containing clearly intact GEMs, representing 28,772 particles, were selected and analyzed in 3D. An *ab initio* generated model from the data by stochastic gradient descent (SGD) was remarkably similar to the previously published (EMDB:9383) *Quasibacillus thermotolerans* iron storage compartment with clear icosahedral symmetry, so this was used for downstream processing. Iterations of homogeneous refinement, local and global CTF refinement and local motion correction yielded a final reconstruction of 2.85 angstroms. The map was post-processed by manually determined B-factor sharpening (B=-100). To retrieve density for the C-terminal GFPs since they represent a breakage of the icosahedral symmetry due to their flexible nature ^48^, homogeneous refinement with pose marginalization and limited alignment was performed. This yielded a reconstruction with a nominal resolution of 4.6 angstroms with clearer density that we ascribed to C-terminal GFP (**Figure 2B**). Measurement of this reconstruction diameter yielded an average of 51 nm.

For 40nm-GEMs, particles were picked with Topaz^49^ yielding a particle set of 29,130 particles. 2D classification was performed on these particles and as done previously, only classes consistent with intact GEM particles were included for further processing. Intact GEM classes represented a particle set of 3,535 particles and again density corresponding to an external cloud of fluorescent proteins was observed. The particles were processed in a similar manner as before with *ab initio* model generation and homogenous refinement with icosahedral symmetry enforced. This yielded a final reconstruction of 4.13 angstroms that was consistent with the biological assembly of the crystal structure of the recombinant *Pyrococcus furiosus* iron storage particle. Since the reconstruction was at a nominally lower resolution than the crystal structure, we opted not to attempt modeling. To resolve the peripheral cloud of fluorescent proteins, we repeated homogeneous refinement with pose marginalization and limited alignment. Single particle workflows are summarized in **Supplementary Figure 3**, map quality metrics and FSC plots are shown in **Supplementary Figure 4** and map and model statistics are summarized in **Table 3**.

#### Model Building of 50nm-GEMs

To model the core scaffold of the 50nm-GEM, the atomic model from the encapsulin iron storage compartment from *Quasibacillus thermotolerans* (PDB:6NJ8) was fit as a rigid body into the 2.85 angstrom map in UCSF ChimeraX ^50^. This was then used as the basis for interactive molecular dynamics flexible fitting (IMDFF) with ISOLDE ^51^. The model was finally refined in Phenix using real space refinement ^52^. Model and map statistics are summarized in **Table 3**. Due to the poor density, we opted not to build an atomic model for the GFP. Structural figures were generated using a combination of UCSF ChimeraX ^50^ and eman2 ^53^.

#### Microscopy of yeast cells

*Saccharomyces cerevisiae* cells were cultured in synthetic complete (SC) medium supplemented with 2% dextrose to mid-log phase (OD_600_ = 0.2–0.4). Prior to imaging, cells were immobilized on 384-well glass-bottom plates coated with Concanavalin A (ConA). Live-cell imaging was performed on a Nikon Ti Eclipse microscope equipped with a 100× oil-immersion objective (NA = 1.4; MRD31901) and an sCMOS camera (Zyla 4.2 Plus, Andor; ZYLA-4.2P-CL10). GFP fluorescence was imaged using an EGFP filter set (EF-EGFP (FITC/Cy2), Chroma) and excitation from a 488 nm laser (OBIS LX 488 nm, 100 mW, Coherent) operated at 100% power. Identical imaging conditions were used for cells expressing 40 nm-GEMs or 50 nm-GEMs. To record GEM dynamics, we used highly inclined thin illumination (HILO) mode. Time-lapse movies were acquired from a single focal plane at a frame interval of 10 ms (100 Hz) with no inter-frame delay, for a total duration of 4 s (400 frames).

#### Microscopy of mammalian cells

HeLa cells grown on tissue-culture treated plates were trypsinized and 0.1×10^6^ cells were plated into each well of a 6 well glass-bottom plate. 24 hours after transfection with GEM expression plasmids, cells were imaged using confocal Nikon TI Eclipse microscope with a spinning disk unit (CSU-X1 spinning disk, Yokogawa, part number = 99459), under a 60x oil objective (60x, NA = 1.49, part number = MRD01691) and a sCMOS camera (Prime 95B, Teledyne Photometrics). The GFP channel was excited through laser light source (OBIS 100mW LX 488 nm, Coherent, part number = 1236444) and imaged through GFP emission filter (EF525/36m, Chroma, part number = 77014803).

#### Mathematical Analysis of Single-Particle Tracking

Let the image sequence be indexed by the discrete set ℱ=1,…,*f*,…,*F*. Each particle trajectory (*i*) is modeled as a discrete stochastic process ***r***_*i*_(*t*) ∈ ℝ^2^, *t* ∈ 𝒯_*i*_⊆ ℱ where 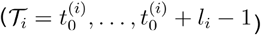, with 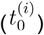 the initial observation time and (*l*_*i*_) the trajectory length. For a lag time (*τ*∈ ℕ), increments of the process are defined asΔ***r***_*i*_(*t; τ*) = ***r***_*i*_(*t+τ*)− ***r***_*i*_(*t*), *t,t* +*τ*∈ 𝒯_*I*_. Assuming weak stationarity of increments, displacement statistics depend only on (*τ*). The empirical propagator ( *p*(Δ***r***; *τ*) was constructed by pooling increments over all trajectories and time origins. For a diffusive process, the propagator is Gaussian

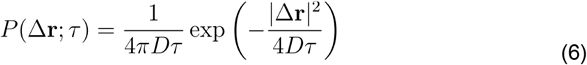

Deviations from Gaussianity were quantified via the non-Gaussian parameter

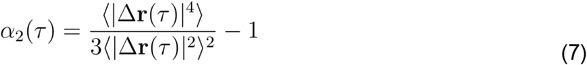

which vanishes for a purely Brownian process. The ensemble mean squared displacement (MSD) is defined as MSD(*τ*)= ⟨|Δ***r*** (*τ*)where the average is taken over particles and time origins. For normal diffusion in two dimensions, MSD(*τ*) = 4*Dτ*. More generally, anomalous transport obeys the scaling law MSD(*τ*) ~ *τ* ^*α*^ with subdiffusion for ( 0<*α*<1) and superdiffusion for (*α*>1). MSD curves were fitted to the generalized diffusion model MSD(*τ*) = 4 *D*_*α*_ *τ*^*α*^ where (*D*_*α*_) is the generalized diffusion coefficient with physical units ( *length*^2^,*time*− ^*α*^). Parameters were estimated by linear regression of (\log(\mathrm{MSD})) versus (\log \tau), restricting the fit to lag times shorter than one quarter of the median trajectory length to reduce finite-track bias. Joint distributions in the parameter space (*D*_*α*_, *α*) were computed across cells; large markers denote per-cell medians, and kernel density estimation provides a nonparametric approximation of the underlying probability density.

#### Further details on analysis of trajectories

The tracking of particles was performed with the Mosaic suite of FIJI^54^, using the following parameters. For yeast 40nm-GEMs and 50nm-GEMs: radius = 3, cutoff = 0, Per/Abs:0.4, a link range of 1, and a maximum displacement of 5 px, assuming Brownian^55^ dynamics. For mammalian cell 40nm-GEMs and 50nm-GEMs: radius = 2, cutoff = 0, Per/Abs:0.2, a link range of 1, and a maximum displacement of 5 px, assuming Brownian dynamics.

Yeast cells were segmented using Cellpose^55^ applied to bright-field images. Mammalian cells were manually segmented using Z-projections of GEMs movies. After segmentation, a mask was saved for each file, which was later used in GEMspa^56^ to segment tracks.

All trajectories were then analyzed with the GEM-Spa (GEM single particle analysis) software package developed in house: https://github.com/liamholtlab/GEMspa/releases/tag/v0.11-beta Mean-square displacement (MSD) was calculated for every 2D trajectory, and trajectories continuously followed for more than 10 time points were used to fit with linear time dependence based on the first 10 time intervals to quantify time-averaged MSD(*τ*)=4*D*_*α*_ *τ*^*α*^, where T is the imaging time interval and *D* is the effective diffusivity with the unit of μm^2^/s. To determine the ensemble-time-averaged mean-square displacement (MSD), all trajectories were fitted with MSD(*τ*)=4*D*_*α*_ *τ*^*α*^ Here *α* indicates diffusion property, with *α* = 1 being Brownian motion, *α* < 1 suggests sub-diffusive motion and *α* > 1 as super-diffusive motion. We used the median value of *D* among all trajectories to represent each condition, and perform normalization to time point 0 min in most of the yeast experiments. For mammalian cell quantification of *D*, the same field of the view of the cytosolic region from individual cell was cropped, and the median value of *D* from all trajectories within the cropped region were quantified.

Step sizes for all trajectories were extracted from GEM-Spa, to reflect the distance (with the unit of μm) of particles traveled for a given time interval: 100 ms for 40nm-GEMs. We next determined both diffusivities and the anomalous exponent of all trajectories by linearizing the anomalous diffusion equation using a logarithmic transformation followed by fitting a linear regression to the transformed data. Figures 3H and I show plots of log_10_ D versus for 40nm-GEMs and 50nm-GEMs respectively.

The next phase involves using Kernel Density Estimation (KDE) to group these data points. The technical details of equation solving and KDE implementation are further explored in the methodology section of the analysis. Rather than using centroids, the resulting medians of the distributions are plotted. This approach is particularly useful during the single-cell analysis, as it provides a clearer depiction of changes in diffusivity.

The diffusion form is estimated from these plots. For Brownian diffusion, a distribution centered around *α* = 1 is expected, with the KDE plot revealing the spread around this central value. This analysis sets the stage for characterizing various motion types, such as macromolecular crowding, which deviates from Brownian motion and displays distinctive patterns. For example, confined motion is often associated with a lower diffusivity.

We analyze the motion of each trajectory by fitting the MSD vs. tau plot to a line. The parameters of this line correspond to the *α* value (slope) and diffusivity (intercept). The one dimensional Einstein equation is ⟨δ*x*^2^⟩ =2*Dτ* ^32^ Which for two dimensions is: MSD(*τ*)=4*Dτ*. To quantify brightness differences, *S. cerevisiae* expressing either construct were imaged on a Nikon Ti TIRF microscope under identical acquisition settings. Mammalian cells were imaged on a Nikon confocal system using matched excitation and detection parameters. Raw images were processed in FIJI. Background was estimated from cell-free regions and subtracted from each image. Regions of interest (ROIs) corresponding to single cells were manually delineated, and the mean pixel intensity within each ROI was extracted. Only cells with particle densities below a predefined threshold—determined through simulations of particle overlap **(Supplementary Figure 1B (Bottom)**)—were analyzed to avoid fluorescence artifacts arising from high particle density. Together, these simulations demonstrate that although higher particle densities can improve sampling under ideal identity preservation, realistic tracking errors rapidly increase identity swaps, ultimately degrading motion analysis. These results define a density-dependent regime for reliable single-particle tracking and justify the density thresholds applied throughout this work.

Extracted intensity values from N = 3 biological replicates and multiple fields of view were pooled and analyzed in Python. Distributions for 40nm-GEMs and 50nm-GEMs were compared using both the Mann–Whitney U test (median differences) and the Kolmogorov–Smirnov test (overall distribution differences).

#### Synthetic Trajectory Simulation and Image Generation

A population of (N) particles was simulated over (F) frames within a three-dimensional rectangular domain ([0,*x*_*max*_] × [0,*y*_*max*_] × [0,*z*_*max*_]). Particle motion was modeled as Brownian diffusion with diffusion coefficient (*D*), identical for all particles and spatial dimensions. At each time step (Δ*t*), particle displacements were drawn independently according to the Einstein diffusion equation^32^ Δ***r***(*t*)=,( Δ *x* Δ *y* Δ *z*), Δ***r***_*i*_~ 𝒩(0,2*D*Δ*t* with uncorrelated displacements in (x), (y), and (z). No explicit boundary conditions were imposed. At each frame, particle positions were projected onto the imaging plane by retaining the lateral coordinates ((x,y)), while the axial coordinate (z) was used to modulate the optical response. Only particles whose projected ((x,y)) coordinates fell within the image boundaries contributed to the rendered image. Synthetic fluorescence images were generated on a (512×512) pixel grid.

The optical response of the imaging system was modeled using a diffraction-limited Airy disk point spread function (PSF) derived from scalar diffraction theory:

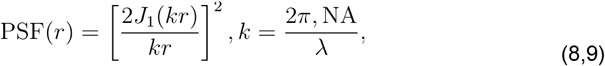

Where (*J*_*i*_) is the first-order Bessel function of the first kind, (*NA*) is the numerical aperture of the objective, and (λ)is the emission wavelength (which we took from microscope parameters. The radial coordinate 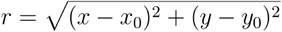 measures the distance from the particle center(*x*_0_,*y*_0_) in the image plane. The PSF was computed on a pixel grid and normalized such that PSF(0)=1. Axial position relative to the focal plane ((*z*_*focus*_=0.5)) modulated both particle brightness and effective PSF width. The PSF radius was scaled quadratically with axial distance:

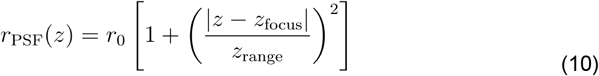

and particle intensity was attenuated according to

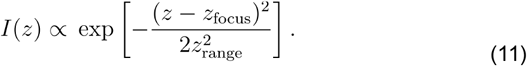

As a result, particles appeared broader and dimmer when displaced from the focal plane.

For each frame, a sparse image was first constructed by placing point emitters at the integer-valued projected particle coordinates. Each emitter was weighted by its axially modulated intensity. The full fluorescence image was then generated by convolving this sparse image with a fixed in-focus Airy disk PSF. To mimic experimental background, additive Gaussian noise was applied independently to each pixel: 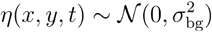, yielding the noisy image *I*_*noisy*_*(x,y,t)*= *I*_*con*_(*x,y,t*)*+ η* (*x,y,t*).

Negative pixel values were clipped to zero: *I*(*x,y,t*) =max! [*I*_*con*_(*x,y,t*),0].

The resulting images were stored as a time-ordered stack, forming a synthetic fluorescence movie. Representative frames were visualized, and the full stack was saved as a multi-frame TIFF file for downstream particle detection and tracking.

#### Error Analysis for simulated trajectories

### Mathematical definition of tracking errors (identity confusion)

Consider a system of (N) particles with true trajectories 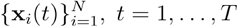 where **x***i*(*t*) ∈ ℝ^2^ denotes the true position of particle (i) at time (t). A tracking algorithm produces a set of reconstructed trajectories 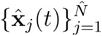 with an associated (possibly time-dependent) assignment map 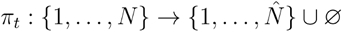 where indicates that true particle is assigned to reconstructed track *j* at time *t*, and *πt*(*i*)= Ø denotes a missed detection. We define an identity-preserving trajectory as one for which the assignment remains constant in time, *πt*(*i*)= *π*_*t*=1_(*i*) ∀*t* An identity error (confusion error) occurs when a particle is either: Missed, i.e.*πt*(*i*)= Ø for at least one *t*, or Swapped, i.e. *πt*(*i*)= ≠ *π*_*t*=1_(*i*) for some (t).

Formally, we define the identity error indicator

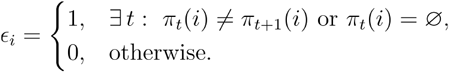

The tracking error fraction is then

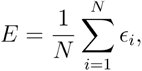

which measures the fraction of particles whose identities cannot be reliably followed over time.

### Density dependence and interpretation

Let *ρ* = *N*/*A* be the particle density in an imaging area *A*. As *ρ* increases, the typical inter-particle distance𝓁~*ρ*^−1/2^ decreases, increasing the probability that two particles come within the tracking algorithm’s spatial or temporal resolution. Identity errors therefore arise not from positional noise, but from ambiguity in particle identity due to spatial overlap, i.e. a loss of distinguishability. In this regime, tracking errors reflect a fundamental information limit imposed by crowding, rather than a failure of the tracking algorithm.

### Fluorescence intensity analysis

Fluorescence intensity was quantified as the mean pixel intensity per particle. The fluorescence intensity of a fluorophore is given by *I* = Φ · *σ* · *I*_*exc*_ where (*σ*) is the fluorescence quantum yield, (Φ) is the absorption cross-section, and (*I*_*exc*_) is the excitation photon flux. For a particle containing (*N*_*f*_) fluorophores, the total emitted intensity is *I*_*total*_ = *N*_*f*_ · Φ ·*σ* · *I*_*exc*_·. Assuming a uniform surface distribution of fluorophores, as expected for GEM assembly, the number of fluorophores scales with particle surface area *N*_*f*_ = *ρ*_*f*_,*S = ρ*_*f*_,4*πR*^2^, Implying *I*_*total*_ *∝ R*^2^ Thus, the expected intensity ratio between 50nm-GEMs and 40nm-GEMs is

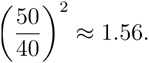

In microscopy images, the emitted light is distributed across multiple pixels according to the point spread function (PSF). Consequently, the intensity measured per pixel depends on both the total fluorescence and the particle’s residence time within the focal plane. For diffusing particles, the measured intensity scales with the time spent in focus, *I*_*measured*_ ∝ *I*_*true*_,*t*_*fouse*_ *·* As a result, mean intensity measurements are systematically biased by particle mobility. In contrast, the maximum pixel intensity, *I*_*max*_ = max(*I*_px_),is less sensitive to focal-plane dwell time and provides a more reliable proxy for fluorophore content. For this reason, all subsequent intensity analyses were performed using (I_{\max}) rather than mean intensity. To validate this approach, we performed simulations of particles undergoing two-dimensional Brownian motion with diffusion coefficient *D* = 0.3μ*m*^2^/*s*, including intermittent transitions to directed motion. These simulations confirmed that mean intensity estimates are increasingly biased with particle mobility, whereas maximum intensity measurements remain stable across dynamical regimes.

#### Software and Algorithms

The following software and algorithms were employed in the analysis: ImageJ, FIJI, Mosaic, Cellpose, GEMspa. The code used in this study is available at https://github.com/liamholtlab

## Materials Availability

Revised atomic model and high resolution core map for the 50nm-GEM were deposited in PDB/EMDB respectively under the accession numbers XXXX/YYYYYY and map for the 40nm-GEM deposited at EMDB under the accession number. For additional information and requests, please contact Liam Holt, Cindy Hernandez, or David C. Duran-Chaparro.

## Acknowledgements

Cryo-EM data for 50nm-GEMs were collected at the Rockefeller University Evelyn Gruss Lipper Cryo-electron Microscopy Resource Center (RRID:SCR_021146) where we thank Mark Ebrahim, Johanna Sotiris and Honkit Ng. Cryo-EM data for 40nm-GEMs was collected at the Simons Electron Microscopy Center at the New York Structural Biology Center, with major support from the Simons Foundation (SF349247) where we thank Jessalyn Miller. This work was supported in part by the National Institutes of Health grants P41GM109824 (MPR) and an Anderson Center for Cancer Research Fellowship at The Rockefeller University (TvE). HeLa cells used in this study were kindly provided by Prof. Jef Boeke (Institute for Systems Genetics, NYU Langone).

## Supplementary Information Movies

**Supp Video 1** | 40nm-GEMs yeast | [Rep1_LH4248_PfVSaph_Basal_001_crop-6.avi](https://drive.google.com/file/d/1OPXkL7H0f8eW3YKuSY_cPjLuTHmDky5r/view?usp=sharing)

**Supp Video 2** | 50nm-GEMs yeast | [Rep1_LH4580_QtESaph_Basal_001_crop-4.avi](https://drive.google.com/file/d/1jLpQWYMQV6xlcTrL9VPB2NmoSZd_z0K-/view?usp=sharing)

**Supp Video 3** | 40nm-GEMs mammalian | [PfV 001-4.avi](https://drive.google.com/file/d/1z7FENiBCX_X_E2f1MZInmRDWeSuUsNQX/view?usp=drive_link)

**Supp Video 4** | 50nm-GEMs mammalian| [QtE 001-4.avi](https://drive.google.com/file/d/1eCe-xXGtVwTl3x9Rq3OywGTXuV-Qc21W/view?usp=drive_link)

**Supp Video 5** | Synthetic Particles n=1 | [syntheticMovie1particle.avi](https://drive.google.com/file/d/1NTQ-ysr_34Qd_WIhnzOTPRLtVoQmFkZm/view?usp=drive_link)

**Supp Video 6** | Synthetic Particles n=10 | [syntheticMovie10particles.avi](https://drive.google.com/file/d/1Oxd8wLlCsW2ms6cEIgzngNO8og8G4sVw/view?usp=drive_link)

**Supp Video 7** | Synthetic Particles n=100| [syntheticMovie100particles.avi](https://drive.google.com/file/d/1AFX4w-u-weB7MBPVG69LNqIGbQha85Pf/view?usp=drive_link)

**Supp Video 8** | Synthetic Particles n=1000| [syntheticMovie1000particles.avi](https://drive.google.com/file/d/1QuVkCsxpqxlyAYl4rhqU93KcQ1i_cc2I/view?usp=sharing)

## Notes

### Competing Interest Statement

The authors have declared no competing interest.

### Summary of Updates

We have added data about experiments in mammalian cells plus simulations of particles to understand the Brownian Motion

